# Distinct genomic and immunologic tumor evolution in germline *TP53-*driven breast cancers

**DOI:** 10.1101/2024.04.03.588009

**Authors:** Nabamita Boruah, David Hoyos, Renyta Moses, Ryan Hausler, Heena Desai, Anh N Le, Madeline Good, Gregory Kelly, Ashvathi Raghavakaimal, Maliha Tayeb, Mohana Narasimhamurthy, Abigail Doucette, Peter Gabriel, Michael J. Feldman, Jinae Park, Miguel Lopez de Rodas, Kurt A. Schalper, Shari B. Goldfarb, Anupma Nayak, Arnold J. Levine, Benjamin D. Greenbaum, Kara N. Maxwell

## Abstract

Pathogenic germline *TP53* alterations cause Li-Fraumeni Syndrome (LFS), and breast cancer is the most common cancer in LFS females. We performed first of its kind multimodal analysis of LFS breast cancer (LFS-BC) compared to sporadic premenopausal BC. Nearly all LFS-BC underwent biallelic loss of *TP53* with no recurrent oncogenic variants except *ERBB2* (HER2) amplification. Compared to sporadic BC, *in situ* and invasive LFS-BC exhibited a high burden of short amplified aneuploid segments (SAAS). Pro-apoptotic p53 target genes *BAX* and *TP53I3* failed to be up-regulated in LFS-BC as was seen in sporadic BC compared to normal breast tissue. LFS-BC had lower CD8+ T-cell infiltration compared to sporadic BC yet higher levels of proliferating cytotoxic T-cells. Within LFS-BC, progression from *in situ* to invasive BC was marked by an increase in chromosomal instability with a decrease in proliferating cytotoxic T-cells. Our study uncovers critical events in mutant p53-driven tumorigenesis in breast tissue.

## Introduction

Acquired, somatic *TP53* alterations are the most common genomic alteration in cancer^1^. Oncogenic alterations in *TP53* lead to the loss of p53 tumor suppressive function, resulting in increased cellular proliferation, genomic instability, and therapy resistance^2,3^. Pathogenic germline variants (PGVs) in *TP53* occur in approximately 1:5000 people and lead to the Li Fraumeni Syndrome (LFS) spectrum in which individuals have an up to 90% risk of developing cancer over their lifetimes^4^. The most common cancer in females with LFS is breast cancer (BC) which occurs at a median age of onset of 30-35, nearly 30 years younger than sporadic BC^5,6^.

Over 70% of sporadic BC are positive for the estrogen receptor (ER+) and negative for receptor tyrosine-protein kinase erbB-2 (HER2)^7,8^. Typically, LFS-BC are ER+; over half are HER2+ and triple negative BC (TNBC) is rarely found in LFS females^6,9^. This is peculiar as somatically acquired *TP53* alterations are present in 80% of sporadic TNBCs versus only 10-15% of sporadic ER+ BC (with or without HER2+)^1^. This distribution of hormone receptor subtypes along with younger age of onset of BC in LFS patients compared to sporadic BC suggests unique mechanisms of breast tumor formation driven by *TP53* PGVs. Indeed, a recent study of pediatric LFS tumors showed that *TP53* PGVs are associated with earlier biallelic loss of *TP53* compared to sporadic *TP53* mutant tumors^10^.

Beyond genomic changes, breast tumor development depends on tumor microenvironment (TME) alterations that facilitate tumor growth. p53 dysfunction may reprogram the TME leading to an altered immunological response which aggravates tumor progression^11,12^. To date, there are no published data analyzing differences in the genomic, transcriptomic, or immune profiles of germline mutant p53 driven BC compared to sporadic BC. The unique characteristics of LFS-BC suggest different tumorigenesis mechanisms compared to sporadic BC that may have implications for early detection and cancer interception in LFS patients. In addition, studying LFS-BC provides a unique opportunity to understand mutant p53-driven breast tumor initiation and development in humans.

## Results

### Clinical characteristics of LFS and early onset non-LFS breast cancer

The Penn Medicine LFS-BC cohort included 93 females with 130 BC cases (**Table 1**); 76% of patients were of self-identified White race/ethnicity, 9% were self-identified Black and 6% were Asian. The pre-menopausal nonLFS-BC cohort included 198 females with 209 breast cancer cases; of these, 69% of patients self-identified as White race/ethnicity, 23% were self-identified Black and 4% were Asian. The median (IQR) age of first BC diagnosis was significantly younger in the LFS-BC compared to the nonLFS-BC cohort [36(29-45) vs 43(39-47), p<0.0001). Fifteen percent of LFS-BC patients were from families who met Classic LFS criteria and 64% met Chompret criteria. LFS-BC patients were significantly more likely than nonLFS-BC patients to have a personal history of breast and another cancer, including another breast cancer (64% vs 18%, p<0.0001). The majority of 74 LFS (92%) and 203 nonLFS (86%) invasive BC were ER+; however, LFS-BC were more likely to be HER2+ (51% vs 20%, p<0.0001). Invasive LFS-BC were more frequently high grade (67% vs 47%, p=0.01). The AJCC stage distribution of LFS-BC and nonLFS-BC were similar.

**Table 1:** Clinical and pathological characteristics of the cohort.

|  | **LFS-BC** |  | **Early onset nonLFS-BC** |  | **p-value** |
| --- | --- | --- | --- | --- | --- |
|  | **N** | **%** | **n** | **%** |  |
| **N/No.BC** | **n=93/130** |  | **n=198/209** |  |  |
| **Age at 1st BC Diagnosis** | **(n=93)** |  | **(n=198)** |  | **<0.0001** |
| Age Median (IQR) | 36 | (29-45) | 43 | (39-47) |  |
| **Self-reported race** | **(n=93)** |  | **(n=198)** |  | **0.0082** |
| Caucasian | 71 | 76.3% | 136 | 68.7% |  |
| African American | 8 | 8.6% | 46 | 23.2% |  |
| Asian | 6 | 6.5% | 7 | 3.5% |  |
| Mixed Race | 2 | 2.2% | 0 | 0.0% |  |
| Other | 6 | 6.5% | 9 | 4.5% |  |
| **LFS classification** | **(n=86)** |  | n/a |  |  |
| Classical | 13 | 15.1% |  |  |  |
| Chompret | 55 | 64.0% |  |  |  |
| LFL (Birch or Eeles) | 48 | 55.8% |  |  |  |
| **Vital status** | **(n=93)** |  | **(n=198)** |  | **0.046** |
| Deceased | 19 | 20.4% | 23 | 11.6% |  |
| **Personal cancer history** | **(n=93)** |  | **(n=198)** |  | **<0.0001** |
| BC + non-breast | 43 | 46.2% | 26 | 13.1% |  |
| Multiple BC | 17 | 18.3% | 10 | 5.1% |  |
| Single BC | 33 | 35.5% | 162 | 81.8% |  |
| **Disease progression** | **(n=91)** |  | **(n=197)** |  | **0.2109** |
| Local recurrence | 4 | 4.4% | 13 | 6.6% |  |
| Distant metastasis | 13 | 14.3% | 37 | 18.8% |  |
| **Breast cancer characteristics** |  |  |  |  |  |
| **Invasive Histology** | **(n=93)** |  | **(n=209)** |  | **0.0015** |
| IDC | 75 | 80.6% | 151 | 72.2% |  |
| ILC and/or Mixed | 10 | 10.8% | 53 | 25.4% |  |
| Other/NOS | 8 | 8.6% | 5 | 2.4% |  |
| **Invasive HR Status** | **(n=74)** |  | **(n=203)** |  | **<0.0001** |
| ER+/HER2- | 30 | 40.5% | 136 | 67.0% |  |
| HER2+ | 38 | 51.4% | 40 | 19.7% |  |
| TNBC | 6 | 8.1% | 27 | 13.3% |  |
| **Grade** | **(n=60)** |  | **(n=171)** |  | **0.0128** |
| Low (1) | 2 | 3.3% | 23 | 13.5% |  |
| Intermediate (2) | 18 | 30.0% | 68 | 39.8% |  |
| High (3) | 40 | 66.7% | 80 | 46.8% |  |
| **Invasive Stage** | **(n=70)** |  | **(n=125)** |  | **0.6538** |
| I | 31 | 44.3% | 45 | 36.0% |  |
| II | 25 | 35.7% | 53 | 42.4% |  |
| III | 11 | 15.7% | 23 | 18.4% |  |
| IV | 3 | 4.3% | 4 | 3.2% |  |

LFL, Li-Fraumeni-like criteria including Birch and Eeles criteria; BC, breast cancer; 1st Dgxn, first diagnosis; Sub. Dgxn, subsequent diagnosis; IDC, invasive ductal carcinoma; DCIS, ductal carcinoma in situ; MDLC, mixed ductal and lobular carcinoma; NOS, not otherwise specified.

### TP53 status of LFS breast cancer

Tumors with sufficient material for p53 IHC and targeted NGS analysis were available for 21 LFS-BC, 15 surrounding normal and 11 contralateral normal specimens (**Figure S1**). Among these, eleven cases had DNA binding domain mutations, including known dominant negative loss of function mutations (G245S, R248Q, R110L) and putative hypomorphic mutations (T125M, R181H, P151S, C141Y), seven had loss of function mutations, and four had tetramerization domain mutations (G334R, R337C, R337H) (**Figure 1a, TableS1**).

**Figure 1:**
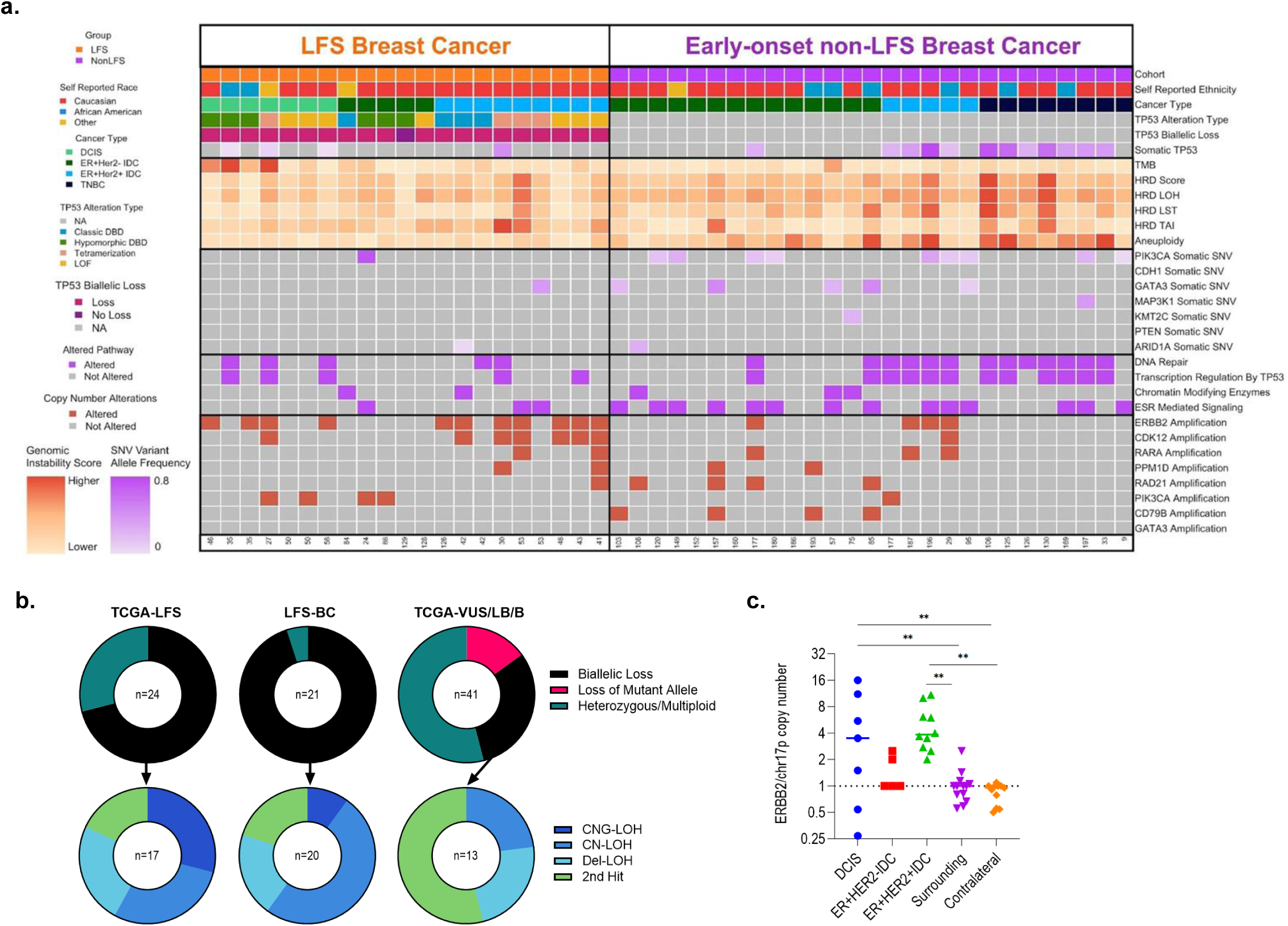
Genomic features of LFS-BC compared to early onset nonLFS-BC. **(a)** Heat-map showing overall clinical and genomic features of LFS-BC and early-onset nonLFS-BC, including self-reported ethnicity, *TP53* germline and somatic variant status and variant alteration type, breast cancer histology and hormone receptor class, *TP53* biallelic loss status, tumor mutational burden, overall homologous recombination deficiency (HRD) score, genomic loss of heterozygosity score, large state transition score, non-telomeric allelic imbalance score, and aneuploidy score, oncogenic mutations, gene-level amplifications/deletions, and mutated gene oncogenic pathways. **(b)** State of the germline *TP53* locus in LFS-BC, tumors from TCGA with germline *TP53* PGVs and tumors from TCGA with germline *TP53* LB/B-VUS. In tumors with biallelic loss, the mechanism of biallelic loss is shown, including copy neutral loss of heterozygosity (CN-LOH), copy neutral gain loss of heterozygosity (CNG-LOH), deletion loss of heterozygosity (Del-LOH), and a second somatic mutation in *TP53* (2^nd^ hit). **(c)** Amplification of *ERBB2* (HER2) in LFS-BC and normal breast tissues. Copy number state for *ERBB2* was normalized to the copy number state of chr17p. Abbreviations: CN, copy neutral; DBD, DNA binding domain; DCIS, ductal carcinoma in situ; Del, deletion; ER, Estrogen Receptor; HRD, homologous recombination deficiency; HER2: receptor tyrosine-protein kinase erbB-2; IDC, invasive ductal carcinoma; LB/B: Likely Benign/Benign variants; LFS, Li Fraumeni Syndrome; LOF, loss of function; LOH, loss of heterozygosity; LP/P, Likely Pathogenic/Pathogenic variants; LST, large state transitions; SNV, single nucleotide variant; TAI, telomeric allelic imbalances; TMB, tumor mutational burden; VUS, variants of uncertain significance.

Sixteen (76%) of LFS-BC had locus-specific LOH and four (19%) had an acquired *TP53* somatic mutation (**Figure 1b, Table S2**). High p53 IHC nuclear expression (>50%) was seen in the majority of LFS-BC from patients with missense mutations, whereas no protein expression (<1%, null pattern) was seen in majority of tumors from carriers of large exonic deletion, truncating, and splicing mutations in *TP53*. Combining p53 IHC and sequencing results, we found evidence of biallelic loss of *TP53* (fully mutant p53) in 20/21 (95%) LFS-BC, 2/15 surrounding normal (13%) and no contralateral normal breast tissue (**Figure 1a**). Four cases (19%, two with T125M and two with R110L) were heterozygous diploid by genomic analysis but had greater than 50% of cells with 3+ p53 IHC staining and were called as biallelic loss, presumably due to LOH not detected by genomic means in tumors with admixture of normal cells.

In order to compare bi-allelic loss rate of LFS-BC to other LFS cancers, we identified germline *TP53* variants in the pan-cancer TCGA cohort^36^. Putative germline variants in *TP53* (VAF>30%) were identified in 69 tumors; 24 were classified as PGVs (TCGA-LFS) and 45 as Variant of Uncertain Significance (VUS) or Likely Benign/Benign (LB/B) variants (TCGA-*TP53*-VUS+LB/B) (**Figure 1b, Table S3**). In the TCGA-LFS cohort, 58% of tumors had LOH and 13% had an acquired *TP53* somatic mutation (71% biallelic loss rate). In contrast, only 32% of cancers with *TP53* VUS/LB/B variants (p<0.001) had LOH. Of tumors with biallelic loss, the mechanism of biallelic loss was similar in both LFS-BC cohorts and TCGA-LFS cohorts with copy neutral LOH being most common (**Figure 1b)**.

### Oncogenic Alterations in LFS versus non-LFS breast cancer

In order to determine whether LFS-BC had any unique oncogenic alterations, we compared LFS-BC to early onset nonLFS-BC from patients negative for PGVs in any cancer risk gene (**Table S4**) and also to nonLFS-BC from TCGA (**Table S5**). Acquired *TP53* oncogenic variants were found in 19% of TCGA-BC ER+Her2-, 37% of HER2+ and 73% TNBC (**Table S5**). Two of 14 (14%) early onset non-LFS ER+Her2-BC had an acquired *TP53* oncogenic variant. The majority of HER2+ (60%) and triple negative (88%) early onset nonLFS-BC had an acquired *TP53* oncogenic variant (**Table S2**).

The TCGA *TP53* mutant or wild-type *TP53* BC had recurrent oncogenic mutations in several genes including *PIK3CA, GATA3, CDH1, MAP3K1, KMT2C* and *PTEN* (**Figure S2a**). In contrast, there were no recurrent oncogenic mutations in LFS-BC (**Figure S2b**). Early-onset nonLFS-BC had recurrent oncogenic mutations in *PIK3CA, GATA3* and *MAP3K1* similar to TCGA; for example, 25-40% of early onset nonLFS-BC depending on subtype versus 4% of LFS-BC had an oncogenic mutation in *PIK3CA* (**Figure 1a, Figure S2c**). We also found no common oncogenic pathways enriched in LFS-BC based on count of mutated genes from the Molecular Signatures Database (MSigDB) (**Figure 1a**). Interestingly, mutations in estrogen receptor (ESR) mediated signaling genes were found in 52% of nonLFS-BC compared to 14% of LFS-BC (p=0.01).

In contrast to infrequent oncogenic mutations, recurrent focal amplifications (normalized CN state ≥4) were seen in LFS-BC, including *ERBB2* (33%) and *CDK12* (33%), (**Figure S2d**). The normalized copy number state of the *ERBB2* locus of all LFS invasive BC was ≥1, all clinically HER2+ LFS-BC had normalized copy number state ≥2 (**Figure 1c**). LFS-DCIS ranged from 0.25-16. Normalized ratios were <1 in several normal LFS breast tissues, but in each case, this was due to amplification of the entire 17p arm with a diploid *ERBB2* locus. Examination of normalized copy number ratios showed that all LFS-BC had co-amplification of *ERBB2* and *CDK12* (**Table S2).**

### Genomic instability in LFS breast cancer

Given p53’s role in maintaining genome integrity, we next compared various measures of genomic instability in LFS-BC to normal tissues. Microsatellite instability (MSI) and tumor mutational burden (TMB) were not significantly different between LFS-BC and normal breast tissues (**Figure 2a**); whereas measures of chromosomal level genomic instability, including whole arm aneuploidy (aneuploidy score), overall chromosomal instability (CIN score), and homologous recombination deficiency (HRD) scores were significantly higher in LFS-BC compared to normal breast tissues. HRD is a composite measure of genomic LOH, large state transitions (LST) and segmental allelic imbalances (nTAI); all of which were increased in LFS-BC compared to normal breast tissue. Indeed, we determined the change in genomic instability measures between pairs of LFS-BC tumors and their normal tissue (**Figure S3a**), showing that only 27% of tumors had higher TMB compared to their matched normal tissue; whereas chromosomal instability measures were uniformly higher in tumors, for example 92% and 100% of tumor-normal pairs showed enrichment of allelic imbalance segments and aneuploid chromosome arms, respectively (**Figure S3b**).

**Figure 2:**
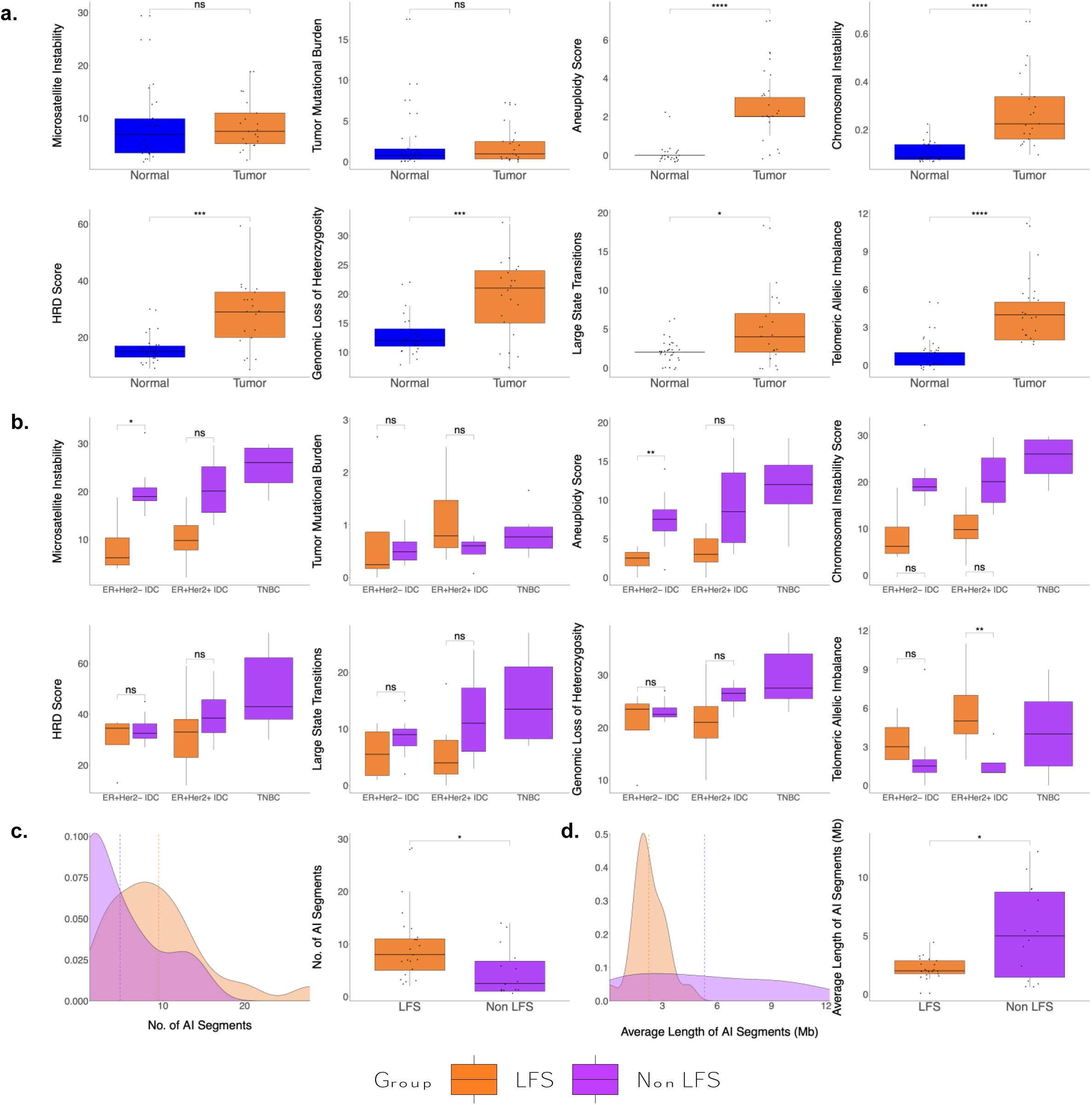
Copy number alterations in LFS-BC compared to early onset nonLFS-BC. **(a)** Genomic instability measures including microsatellite instability (MSI), tumor mutational burden (TMB), aneuploidy score (AS), chromosomal instability (CIN) score, homologous recombination deficiency (HRD) score, genomic loss of heterozygosity (gLOH), large state transitions (LST) and non-telomeric allelic imbalances (TAI) in normal LFS breast tissue versus LFS-BC. **(b)** Genomic instability measures (MSI, TMB, CIN, HRD, gLOH, LST, nTAI) stratified by hormone receptor status in LFS-BC versus nonLFS-BC. **(c)** Distribution of the number of amplified (total copy number ≥5) segments of allelic imbalance, aka short amplified aneuploid segments (SAAS) across LFS-BC and nonLFS-BC and box plot of number of amplified AI segments in LFS-BC versus nonLFS-BC. **(d)** Distribution of the lengths of amplified AI segments in LFS-BC versus nonLFS-BC and box plot of average length of amplified AI segments in LFS-BC versus nonLFS-BC. Abbreviations: AI, allelic imbalance; Avg, average; CNA, copy number alteration; CNt, copy number total. ER, Estrogen Receptor; HRD, homologous recombination deficiency; HER2: receptor tyrosine-protein kinase erbB-2; No, number; IDC, invasive ductal carcinoma; LFS, Li Fraumeni Syndrome; TNBC, triple negative breast cancer. *p<0.05; **p<0.01; p<0.001; ***p<0.0001, uncorrected.

We next determined whether these genomic instability measures differed between LFS-BC and early onset nonLFS-BC by hormone receptor subtype. All genomic instability measures showed the expected trend of increased levels of instability in triple negative versus ER+ nonLFSBC (**Figure 2b**). Interestingly, nearly all genomic instability measures were similar or lower in LFS-BC compared to early onset nonLFS-BC. In ER+ BC, MSI and aneuploidy score were significantly lower in LFS-BC versus nonLFS-BC, and CIN, HRD and its components were similar across hormone receptor (HR) subtypes between LFS-BC and nonLFS-BC. Only segmental allelic imbalances were higher in LFS-BC, particularly HER2+ subtypes, compared to nonLFS-BC. The per tumor average number of segments of allelic imbalance (AI) with at least five-fold amplification were significantly higher in LFS-BC versus nonLFS-BC (**Figure 2c**). Interestingly, these aneuploid segments were significantly shorter in LFS-BC than nonLFS-BC (**Figure 2d**). and included many oncogenes such as *ERBB2* (**Figure S2**). In contrast, in sporadic BC from TCGA, aneuploidy score and all HRD measures were higher across all hormone receptor subtypes in BC with acquired *TP53* mutations compared to *TP53*-WT tumors (**Figure S4**), demonstrating that enrichment of all levels of genomic instability occurs with acquired p53 mutations; however, there is a specific enrichment of only short aneuploid amplified segments (SAAS) in LFS-BC tumors.

In order to further explore the copy number signatures unique to LFS-BC, we performed whole genome sequencing of LFS DCIS (n=5), ER+Her2-BC (n=4) and HER2+ BC (n=10) (**Figure S5**). Consistent with the enrichment of SAAS, the most prominent copy number signatures in LFS-BC were chromosomal instability and chromothripsis (CN9/CN5); only two LFS-BC showed evidence of whole genome doubling and diploidy.

### Mutational processes in LFS breast cancer

Having demonstrated the predominance of SAAS in LFS-BC, we next examined the single nucleotide substitution mutational processes seen in LFS-BC (**Figure S5**). All samples had SBS1 and 75% had SBS5 signatures (clock-like signature associated with spontaneous 5-methylcytosine deamination and unknown etiology, respectively). In addition, nearly all LFS-BC had SBS3 (HRD, 10/12) and SBS18 (ROS, 11/12). Other BC associated signatures, SBS2 and SBS13 (APOBEC activity), were not seen in LFS-BC.

### Gene expression patterns in LFS breast cancer

To examine whether LFS-BC demonstrates a pattern of gene expression reflective of p53 functional loss, we performed differential expression analysis on bulk RNA-seq from LFS-BC versus LFS normal breast tissues and compared this to differential expression analysis of nonLFS-BC versus non-LFS normal breast tissues from the early onset nonLFS-BC cohort and TCGA-BC, stratified by hormone receptor subtype and somatic *TP53* mutation status. Principal component analysis (PCA) plots of LFS-BC RNAseq data demonstrated clustering of normal (contralateral and surrounding) and tumor tissue (**Figure S6a**) and clustering of hormone receptor subtypes (**Figure S6b**); in addition, no batch effects were seen between LFS-BC and nonLFS-BC data except for nonLFS-TNBC (**Figure S6c-d**) which were then excluded from downstream differential expression comparisons. We observed 2352 significantly differentially expressed genes (DEGs) in LFS-BC compared to LFS normal breast tissues (adjusted p-value<0.05 and log2fold change >+/-1.5) (**Figure 3a, Table S6**). Similarly, we observed 2547 DEGs in nonLFS-BC compared to nonLFS normal breast (**Figure 3b, Table S7**). More DEGs were identified in TCGA-BC hormone receptor subtypes compared to normal breast tissue as expected with larger numbers of samples, but similar numbers of DEGs were identified regardless of somatic *TP53* mutation status (**Figure S7, Table S8-S13).**

**Figure 3:**
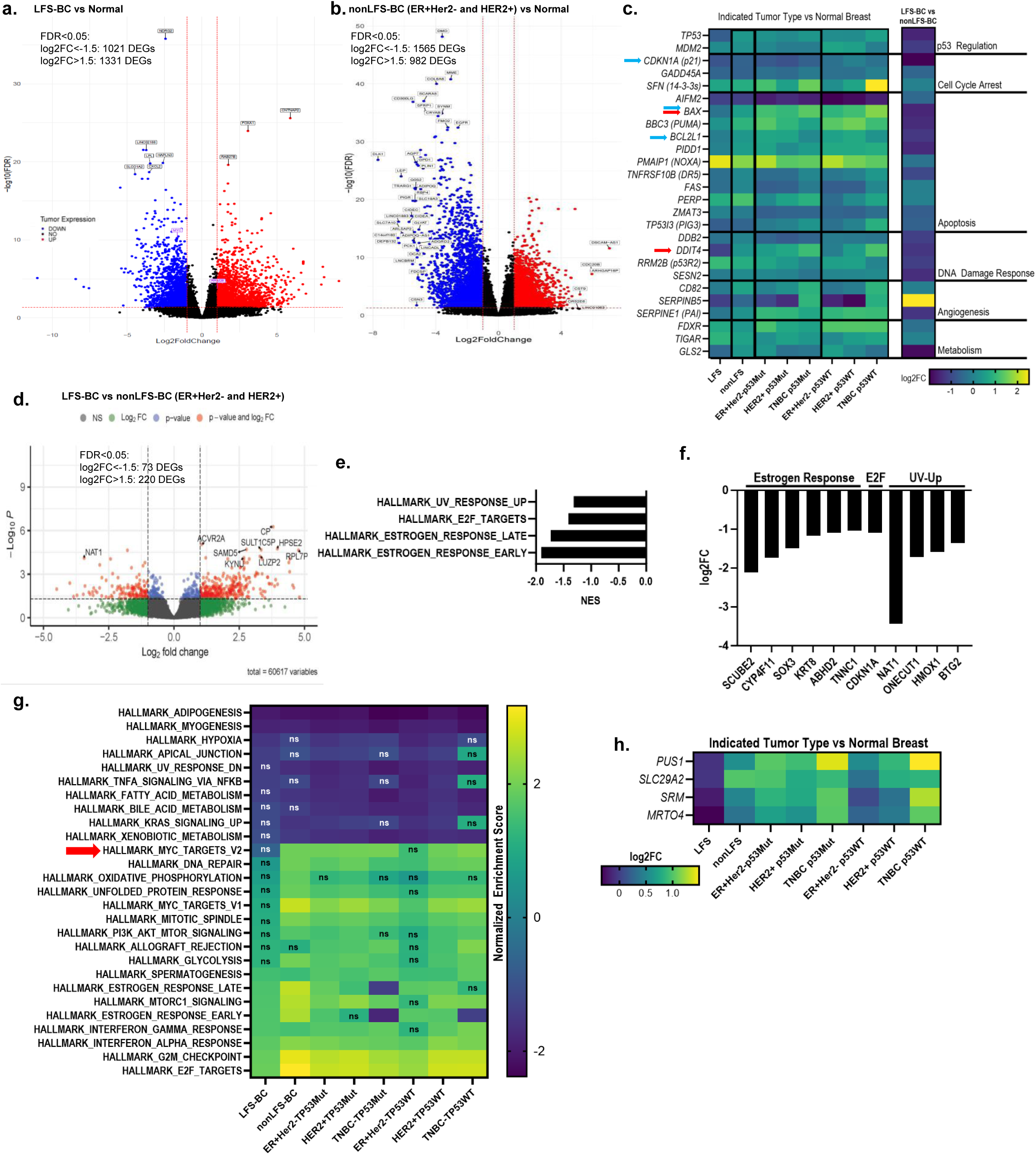
Gene expression profiling in LFS-BC compared to early onset nonLFS-BC. **(a)** Volcano plot showing differentially expressed genes (DEGs) in LFS-BC compared to normal LFS breast tissue. **(b)** Volcano plot showing DEGs in nonLFS-BC compared to normal breast tissue. **(c)** Differential expression of canonical p53 targets in the indicated tumor types (LFS-BC, early onset nonLFS-BC, TCGA ER+Her2-nonLFS-BC stratified by somatic *TP53* mutation status, TCGA HER2+BC nonLFS-BC stratified by somatic *TP53* mutation status, and TCGA non-LFS TNBC stratified by somatic *TP53* mutation status) compared to normal breast tissue. **(d)** Volcano plot showing DEGs in LFS-BC compared to early-onset nonLFS-BC. **(e)** Hallmark pathways significantly differentially regulated between LFS-BC and nonLFS-BC. **(f)** Log2 fold change (FC) of significantly regulated genes in these pathways in LFS-BC compared to nonLFS-BC. **(g)** Normalized enrichment scores for Hallmark pathways in differential gene expression analyses between LFS-BC, nonLFS-BC and TCGA ER+, Her2+ and TNBC, each stratified by somatic p53 mutation status and each group compared to breast tissue normal. Pathways shown were significantly enriched in at least one comparison, non-significantly enriched pathways are indicated by “ns”; red arrow indicates pathway significantly regulated in all comparisons except LFS-BC versus normal. **(h)** Log2 fold change (FC) of significantly regulated genes in MYC targets pathway in indicated tumor type compared to normal breast tissue. Abbreviations: BC, breast cancer; DEG, differentially expressed genes; ER, Estrogen receptor; FC, fold change; FDR, false discovery rate; HER2: receptor tyrosine-protein kinase erbB-2; LFS, Li Fraumeni Syndrome; p53Mut, positive for somatic *TP53* mutation; p53WT, negative for somatic *TP53* mutation; TNBC, triple negative breast cancer.

We next examined differential expression of a well-curated list of 326 p53 target genes^44^ in LFS-BC, nonLFS-BC and each TCGA subset compared to normal breast tissue, hypothesizing that *TP53* target genes important for BC development would be significantly up-regulated in nonLFS-BC and TCGA-BC versus normal breast tissue but not regulated or down-regulated in LFS-BC versus normal breast tissue. Most p53 target genes were regulated in similar directions and magnitude; notably, however, *BAX* and *DDIT4* were significantly up-regulated in all nonLFS-BC cohorts versus normal breast tissues but either down-regulated or not regulated in LFS-BC versus normal breast tissue (**Figure 3c**, red arrows). We next examined these genes in a comparison of LFS-BC versus nonLFS-BC tumors (**Figure 3c, Table S14**). Expression of p53 apoptosis targets *BAX* and *BCL2L1* and cell cycle arrest gene *CDKN1A* (p21) were significantly lower in LFS-BC versus nonLFS-BC (blue arrows).

Overall, comparing LFS-BC to nonLFS-BC in a tumor-to-tumor comparison, we found 73 genes that were significantly down-regulated and 220 significantly up-regulated in LFS-BC versus nonLFS-BC (**Figure 3d**). MSigDB Gene set enrichment analysis (GSEA), identified estrogen response, E2F and UV response as the most significant hallmark pathways that were negatively regulated in LFS-BC vs nonLFS-BC (**Figure 3e, Table S15**). The genes that were significantly down-regulated in LFS-BC versus nonLFS-BC in these pathways included tumor suppressors such as *CDKN1A* and *BTG2*. (**Figure 3f**). We next performed GSEA on the LFS-BC, nonLFS-BC and TCGA cohort BC versus normal breast tissues (**Figure S8, Table S16**). Most significantly up and down regulated Hallmark pathways in LFS-BC were similarly regulated in nonLFS-BC and all TCGA-BC (ER+Her2-and HER2+) subtypes (**Figure 3g, Figure S8**). Notably, however, MYC pathway was significantly up-regulated in most nonLFS-BC and TCGA cohorts and not up-regulated in LFS-BC, four genes from this pathway were significantly downregulated in LFS-BC versus normal breast tissue whereas those genes were significantly upregulated in non-LFS BC versus normal (**Figure 3h**).

### Tumor microenvironment in LFS breast cancers

In BC, the presence and activity of tumor-infiltrating lymphocytes have significant implications for the prognosis and treatment^45^. We therefore investigated the tumor immune microenvironment (TIME) in LFS-BC compared to nonLFS-BC and the normal breast tissues from each group using the RNASeq data. CIBERSORT (**Table S17**) was used to determine differences in the proportion of immune cell populations. MCPCounter (**Table S18**) and xCell (**Table S19**) were used to estimate relative abundances of immune cell populations. We observed prominent variation within each group for all TIME cell types (**Figure S9**). We observed no differences in composite stroma, microenvironment or immune scores in invasive LFS-BC compared to early onset nonLFS-BC (**Figure 4a**).

**Figure 4:**
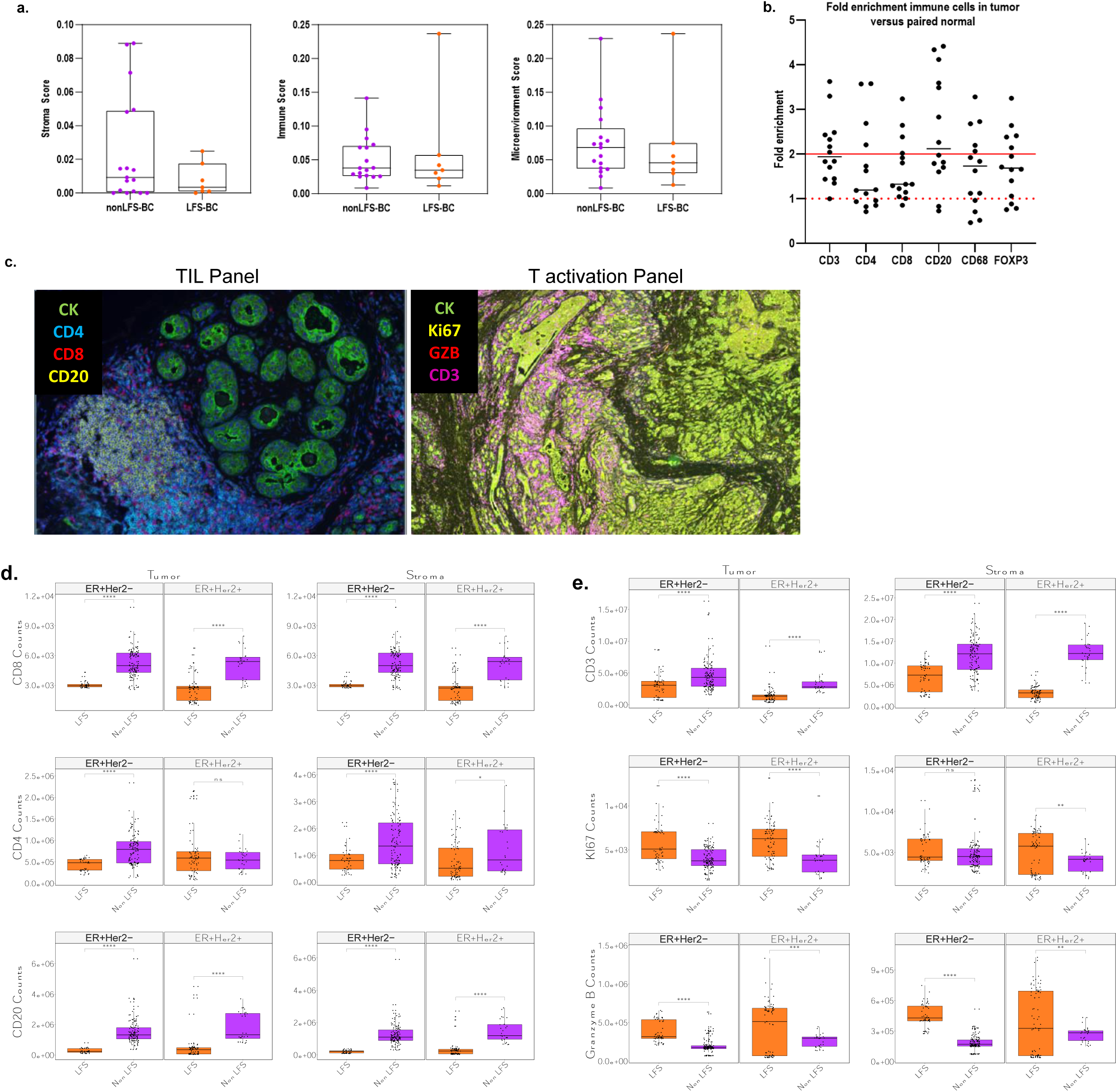
Immunological profiling in LFS-BC compared to early onset nonLFS-BC. **(a)** xCell analysis of RNASeq data showing stroma, immune and microenvironment scores from invasive cancers from LFS-BC and early-onset nonLFS.. **(b)** Single stain IHC in LFS-BC compared to normal LFS breast tissue, fold enrichments in tumor versus normal are shown. **(c)** Representative images of multiplex immunohistochemistry (IHC) results staining for tumor infiltrating lymphocyte (TIL) and T-cell activation (ACT) panels. **(d)** Quantification of multiplex IHC results for TIL panel in LFS-BC, nonLFS ER+BC, nonLFS-TNBC. CD4, CD8 and CD20 in tumor and stroma are shown. Abbreviations: ACT, T-cell activation; CK, cytokeratin; BC, breast cancer; ER, estrogen receptor; HER2: receptor tyrosine-protein kinase erbB-2; LFS, Li Fraumeni Syndrome; TIL, tumor infiltrating lymphocytes; TNBC, triple negative breast cancer.

We next performed single immune cell marker IHC on the LFS-BC compared to surrounding normal tissue. Comparing an LFS breast tumor to its surrounding normal demonstrated that CD3+, FoxP3+ Tregs, CD68+ and CD20+ cells were approximately two-fold enriched in the majority of tumor-normal pairs compared to surrounding normal breast tissue (**Figure 4b**).

We assayed our LFS-BC versus early-onset nonLFS-BC samples using two mIF panels which tested the levels of specific immune cell subpopulations (TIL, ACT) (**Figure 4c**). The levels of the markers were selectively measured in three different spatial tumor tissue compartments using fluorescence co-localization strategies^40^ including: the cancer-cell nests (CK+ pixels/area or tumor compartment), the non-tumor/stromal cells (CK-pixels/area or stromal compartment) and the entire tumor tissue including both malignant and non-malignant cells (dilated DAPI+ pixels/area).

We first examined whether our method recapitulated a difference in immune cell presence in non-LFS. We found early onset non-LFS TNBC had significantly higher densities of CD8+, CD4+, and CD20+ cells in their stroma compared to ER+Her2- and HER2+ nonLFS-BC with similar levels intratumorally (**Figure S10a-c**). We discovered that compared to early onset nonLFS-BC, LFS-BC had significantly lower tumor infiltrating CD8+ and CD20+ cells in both HER2- and HER2+ cancers (**Figure 4d**). In addition, CD4+ TIL levels were also lower in LFS-BC than non-LFS but only in HER2-tumors. These differences occurred throughout compartments (**Figure S11a**).

We next assayed for T-cell activation using CD3, granzymeB and Ki-67. As expected, CD3 and granzyme B levels were correlated and Ki67 was higher in tumor than stroma (**Figure S11b**). While LFS-BC had lower densities of CD3+ cells compared to early-onset nonLFS-BC, LFS-BC had significantly higher levels of granzymeB and Ki-67 (**Figure 4e**), indicating higher levels of activated cytotoxic T-cells in LFS-BC. This was found in both HER2- and HER2+ comparisons and across compartments (**Figure S11c**).

In order to validate these results in a larger cohort, we additionally evaluated 17 LFS-BC and 24 nonLFS-BC from MSKCC. The MSKCC LFS-BC cohort was older and tumors were lower stage compared to the Penn LFS-BC cohort; the distribution of hormone receptor status and grade were similar (**Table S20**). The nonLFS-BC cohorts had similar clinical and pathological features (**Table S20**). Combining these cohorts recapitulated the findings of lower CD8+ and CD20+ cells and higher levels of Ki67 and granzymeB in LFS-BC versus nonLFS-BC tumors (**Figure S12a-d**).

### Evolution of genomic, transcriptomic and immune cell changes in DCIS compared to invasive LFS breast cancer

Invasive BC are thought to arise through a phase of DCIS; therefore, we examined the genomic, transcriptomic and immune cell differences between DCIS and IDC in LFS-BC. When LFS-BC were stratified into DCIS versus IDC, all chromosomal level genomic instability measures, except LSTs, were statistically higher in both DCIS and IDC compared to normal tissue. Genomic instability was similar although trended towards an increase in the IDC versus DCIS tumors (**Figure 5a**). LFS DCIS and IDC had significant differences in gene expression profile (**Table S20**). We observed 275 differentially expressed genes in LFS-IDC compared to LFS-DCIS (**Figure 5b**); with a number of oncogenic pathways significantly up-regulated in LFS-IDC compared to LFS-DCIS (**Figure 5b, Table S21**).

**Figure 5:**
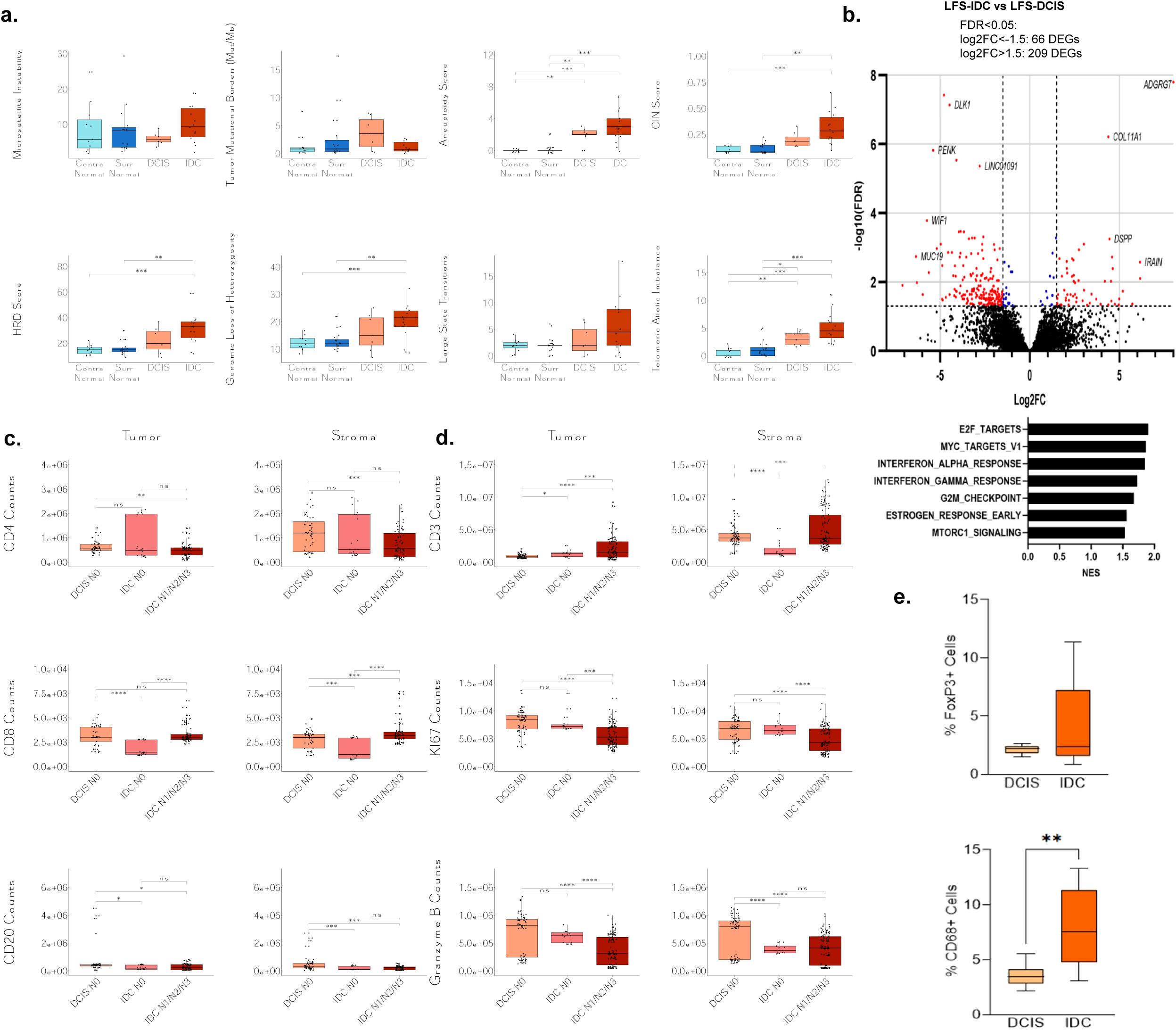
Genomic, transcriptomic and immunological features of LFS-DCIS versus LFS-IDC. **(a)** Genomic instability measures including microsatellite instability (MSI), tumor mutational burden (TMB), aneuploidy score (AS), chromosomal instability (CIN) score, homologous recombination deficiency (HRD) score, genomic loss of heterozygosity (gLOH), large state transitions (LST) and non-telomeric allelic imbalances (TAI) in LFS-DCIS compared to LFS-IDC**. (b)** Differentially expressed genes (DEGs) in LFS-DCIS versus normal LFS breast tissue and LFS-IDC versus normal LFS tissue. Involvement of these DEGs in Hallmark biological processes. **(c)** Quantification of multiplex immunohistochemistry (IHC) results of tumor infiltrating lymphocyte (TIL) panel components in LFS DCIS versus LFS-IDC. CD4, CD8, and CD20 in tumor are shown. **(d)** Quantification of multiplex IHC ACT panel components in LFS-DCIS versus LFS-IDC. Granzyme B, Ki67, and CD3 in tumor are shown. **(e)** Quantification of single stain IHC for FoxP3+ and CD68+ cells in LFS-DCIS versus LFS-IDC. Abbreviations: ACT, T-cell activation; CK, cytokeratin; DCIS, ductal carcinoma in situ; ER, estrogen receptor; Grzmb, granzyme B; HER2: receptor tyrosine-protein kinase erbB-2; IDC, invasive ductal carcinoma; LFS, Li Fraumeni Syndrome; TIL, tumor infiltrating lymphocytes; TNBC, triple negative breast cancer.

When we compared the TIL panel in LFS DCIS versus LFS IDC stratified into tumors that were node negative versus node positive, there were no consistent differences in T or B-cells (**Figure 5c**). LFS-DCIS had significantly higher CD8+ T cell levels compared to node negative IDC but not node positive (**Figure 5c**). These results were recapitulated with single immune marker IHC for CD3, CD4, CD8, and CD20 in a subset of samples (**Figure S13a**), and results were similar in the DAPI compartment (**Figure S13b**). We next quantified the activity of the infiltrating immune compartments using the ACT activity assay in LFS-DCIS versus LFS-IDC. We found that there were significantly higher Ki67+ and Granzyme B in LFS-DCIS as compared to LFS-IDC but fewer CD3+ cells (**Figure 5d**). Furthermore, single immune marker staining in a subset of samples demonstrated a nominal increase in tumor supportive FoxP3+ T-regulatory cells and a significant increase in CD68+ macrophages in LFS-DCIS versus LFS-IDC (**Figure 5e**). These findings suggest that LFS-IDC develop mechanisms for adaptive immune evasion during progression.

## Discussion

In this study, we leveraged patients with germline *TP53* PGVs to identify the unique genomic, transcriptomic and immunologic characteristics of breast tumor formation induced by mutant p53. Breast tumors in LFS patients uniformly underwent biallelic loss of *TP53*, and compared to age-matched early onset non-genetically driven BC, rarely accumulated other oncogenic mutations, and instead accumulated short amplified aneuploid segments, which we term “SAAS”. LFS breast tumors specifically lost transactivation of apoptosis genes versus other p53 target genes when compared to normal breast tissue. Further, LFS-BC had a unique immune infiltrate with overall lower CD8+ T-cells, but higher activated cytotoxic T-cells. Finally, progression of *TP53* mutant *in situ* to invasive tumors was not associated with significant genomic changes but was associated with reduced TILs, loss of cytolytic T cells and prominent upregulation of CD68+ tumor-associated macrophages.

Our data suggest that *TP53* related breast tumor formation in the germline state requires biallelic loss of *TP53* as a first step in tumor formation. This is consistent with data from pediatric tumors in LFS patients^10^ which showed that *TP53* biallelic loss occurs early in tumor formation. Our comprehensive analysis of oncogenic mutations, copy number changes and mutational signatures further shows that biallelic *TP53* loss does not lead to accumulation of single nucleotide variants nor is marked by specific single base pair substitution signatures, likely explaining the absence of other oncogenic mutations in LFS-BC. Instead, we observed that while all levels of genomic instability accumulated in LFS-BC compared to normal LFS breast tissue, only SAAS events were higher in LFS-BC compared to nonLFS-BC. This contrasts with somatic *TP53* mutations which are associated with increases in all levels of genomic instability compared to BC without acquired *TP53* mutations. Other types of genomic level alterations such as genome doublings were not observed. Taken together, mutant p53 driven SAAS formation is likely responsible for the observation that HER2 amplification is a common feature of LFS-BC.

In contrast to our results in BC, a pancreatic cancer *TP53* mutant model demonstrated that *TP53* LOH led first to deletion events followed by genome doubling and amplifications^46^. We found no evidence of genome doubling by copy number signature analysis in LFS-BC. It is known that estrogen stimulates DNA double strand breaks (DSB) in cells as part of its metabolism and proliferative effect^47^ and can lead to translocation-bridge (TB) mediated focal amplifications^48^. Expression profiling in our analysis showed that LFS-BC failed to upregulate estrogen responsive pathways as occurs in nonLFS-BC. Therefore, it is possible that p53 dysfunction leads to a different program of genome instability in estrogen responsive tissues compared to other tissues. Other groups have shown that homologous recombination deficiency (HRD) occurs in *TP53* mutated samples irrespective of HR gene alterations^49^; however, we did not observe higher HRD in LFS-BC compared to nonLFS-BC and DNA damage response pathway were not dysregulated. This again shows that tumorigenesis follows distinct, likely tissue-specific, pathways when p53 dysfunction is the initiating event versus a later event in tumor evolution.

Given a lack of other oncogenic processes, outside of reactivating p53 function in breast tissues, genomic sequencing alone does not suggest clear pathways for specific targeted treatment. However, our previous work on modeling mutant *TP53* fitness predicted selective pressure from T-cells may exist in LFS due to intrinsic trade-offs, with the selective benefit mutant p53 provides cancer cells, implying immunotherapeutic interventions may be possible^50^. Indeed, TNBC which typically has mutant p53 is partially sensitive to immune checkpoint blockade^51^. Unfortunately, we found that LFS-BC had lower CD8+ T-cells than nonLFS-BC, a finding that is consistent with prior studies showing reduced TILs in LFS-BC^9,52^, likely due to higher levels of aneuploidy in LFS-BC^53^. It is intriguing that while CD8+ cells were found at lower levels in LFS-BC, markers of T-cell activation were higher, suggesting possible benefit for an anticancer vaccine to boost tumor specific antigen responses.

There are limitations to our data in that while the largest described cohort of LFS-BC from a genomics and immunological perspective, this sample set is still small limiting comparisons within hormone receptor and stage subtypes. In addition, genomic analysis is limited from FFPE tissues, and additional studies, particularly at the single cell level from fresh tissues will be necessary to confirm these findings and explore the cell intrinsic and extrinsic mechanisms of tumor formation initiated by p53 loss.

In conclusion, breast cancer is the most common cancer in females with LFS, occurring 30 years earlier than sporadic breast cancers. Our data suggest that breast tissues with a germline*TP53* pathogenic variant first undergo biallelic loss of *TP53* to a fully mutant p53 state, resulting in accumulation of SAAS, possibly selecting for *ERBB2* amplification. Mutant p53 leads to transcriptional reprogramming away from an apoptotic state. As LFS-BC progress from DCIS to IDC, a tumor-promoting immune evasive or tolerogenic environment is established. Our study represents the first complete portrait of human p53-driven breast tumors and suggests unique mechanisms of treatment.

## Online Methods

### Cohort ascertainment

Patients with *TP53* PGVs were ascertained from the Penn/CHOP Li Fraumeni Syndrome/TP53 Biobank (NCT04367246). Acquisition of patient blood and tumor samples was approved by the Institutional Review Boards of the University of Pennsylvania and Children’s Hospital of Philadelphia (CHOP) (Penn IRB#834147/CHOP IRB#18-015810). Informed consent was obtained from each participant for use of their samples and clinical data in genetic and immunologic studies. Inclusion criteria for LFS-BC were 1) clinically confirmed *TP53* PGV; 2) available pathology slides and/or tumor blocks; and for sequencing 3) available germline blood DNA. Only primary breast tumors were included. Chart review was performed via the associated research database and the Penn Medicine electronic health record for clinical characteristics, tumor information, and treatment information. Ninety-three LFS-BC patients with 130 primary breast tumors were identified in the Penn Medicine cohort. Sixty-one cases had available slides for review, 56 cases (43%) had identifiable material and a matched germline blood specimen. Specimens from twenty-six LFS-BC (46%) from 19 patients were obtained, of which, surrounding normal breast tissue (SNBT) and contralateral normal breast tissue (CNBT) were available for analysis in 15 cases and 11 cases respectively (**Figure S1, Table S1**).

An early-onset nonLFS-BC cohort was created from Penn Medicine Biobank (PMBB)^13,14^. Acquisition of patient blood and tumor samples was approved by the Institutional Review Board of the University of Pennsylvania (Penn IRB#832122). Patients were included if they had 1) BC diagnosed age<50, 2) undergone germline whole exome sequencing (WES) and had no identified PGV in *TP53* or any of 44 other autosomal dominant cancer susceptibility genes (**Table S4**); and 3) cancer clinical data present in the Penn Medicine Cancer Registry (PMCR). One hundred ninety-eight early onset non-germline genetic driven BC patients with 209 primary breast tumors were identified in the PMBB cohort. Seventy-four cases (37%) had not received pre-operative chemotherapy, had identifiable pathological specimens and a matched germline blood sample. Twenty-nine breast tumors (39%) from 26 patients were obtained for analysis (**Figure S1, Table S1**).

Tissue and plasma samples from LFS-BC and nonLFS-BC at Memorial Sloan Kettering Cancer Center (MSKCC) were identified for validation cohort. These tumors were examined by a trained pathologist team, under research protocols 12-245 or 06-107. Inclusion criteria for the LFS-BC cohort were individuals with germline mutations in *TP53* with a diagnosis of breast cancer. Inclusion criteria for the nonLFS-BC cohort were individuals without germline *TP53* mutations with a diagnosis of breast cancer.

For both cohorts, positivity for estrogen receptor (ER), progesterone receptor (PR) and erbb-2 (HER2) were obtained from clinical pathological reports which follow College of American Pathologists (CAP) reporting^15,16^.

### Sample preparation of sequencing libraries

For the Penn Medicine cohorts, formalin-fixed paraffin-embedded (FFPE) tumor blocks from LFS and nonLFS-BC cases were sectioned and stained with hematoxylin and eosin (H&E) to ensure that sections of over 70% of *in-situ* or invasive tumor were used for DNA and RNA extraction. Selected tumor areas of slides were macro-dissected and prepared for sequencing as previously described^17^. Contralateral and surrounding normal tissue sections were stained with H&E to ensure sections of normal tissue were used for DNA and RNA extraction. Germline DNA from blood or saliva was extracted using standard protocols. Libraries of tumor, normal tissue and matched germline DNA were prepared using KAPA Hyper Prep Kit (Roche Diagnostics, Branchburg NJ). Libraries were subjected to targeted next generation sequencing (NGS) using a custom targeted hybrid capture based probe set including 512 cancer genes^18^ along with the xGen CNV Backbone Panel using xGen Hybridization Protocol (Integrated DNA Technologies, Coralville, IA). Tumors were sequenced on an Illumina Nova-Seq (Illumina, Madison, WI). LFS breast tumors were additionally subjected to whole genome sequencing on an Illumina Nova-Seq (Illumina, Madison, WI). RNA from LFS-BC, surrounding and contralateral normal LFS breast, nonLFS-BC and surrounding normal non-LFS breast samples were prepared using the Stranded Total RNA Prep with Ribo-Zero Plus (Illumina, Madison, WI), creating cDNA libraries. cDNA libraries were sequenced on an Illumina Nova-Seq (Illumina, Madison, WI). For whole genome sequencing, LFS-BC tumor DNA underwent Library Preparation EF 2.0 with Enzymatic Fragmentation and Twist Universal Adapter System (Twist Bioscience, San Francisco, CA).

### Bioinformatics for Targeted Next Generation Sequencing data

FASTQ files from sequencing of 22 LFS-BC, 26 LFS normal breast tissues, 26 nonLFS-BC, and patient-matched blood or saliva normal were aligned to human genome version 19 (hg19) using BWA-mem version 0.7.17^19^. Sequencing quality control removed one LFS-BC and one LFS surrounding normal tissue for a total of 21 LFS-BC, 15 surrounding normal LFS breast tissue, 11 contralateral normal LFS breast tissue, and 26 nonLFS-BC with targeted NGS data **(Table S1)**. Germline variants were called from BAM files using Genome Analysis Toolkits (GATK) HaplotypeCaller version 3.7^20^. Individual tumor sequencing data and normal breast tissue sequencing data were matched to sequencing data from the same patient’s lymphocytes. Somatic copy number variations were called using both Sequenza version 3.0.0^21^ and CNVKit version 0.9.9^22^. Using CNVKit copy number segments as input, HRDex version 0.0.0.9^23^ was used to determine homologous recombination deficiency (HRD) scores including segments of large state transitions (LSTs), genomic loss of heterozygosity (LOH) and non-telomeric allelic imbalance (nTAI) and aneuploidy scores. Copy number burden as a measure of overall chromosomal instability (CIN) was calculated as the percentage of bases measured that were in an altered copy number state. Somatic single nucleotide variants (SNVs) were called using Mutect2 version 4.1.2^24^. Somatic SNVs were annotated using ANNOVAR version 2018-04-16^25^ and OncoKB^26^. Tumor mutational burden (TMB) was calculated as the number of non-oncogenic SNVs with a depth greater than 20 and an allelic balance greater than 0.05 divided by the number of bases captured and multiplied by 100000. Microsatellite instability (MSI) scores were calculated using MSISensor version 0.6^27^..

### Determination of locus-specific loss of heterozygosity for TP53

The presence of genomic locus-specific LOH in *TP53* was determined for each LFS-BC and normal breast tissue samples using a combination of somatic variant allele frequency (VAF) of the PGV as determined by Varscan2, tumor purity as determined by Sequenza, and copy number at the genomic locus as determined by Sequenza. To correct for tumor purity, a minimum expected LOH VAF was calculated as a function of the tumor purity: E = P + (1-P)/2, where E = Expected LOH VAF of 50% and P = Tumor purity. If the observed somatic VAF exceeded the expected LOH VAF and/or the ‘B allele’ copy number was zero, the variant was classified as having locus-specific LOH. The presence of second somatic hits in *TP53*, depending on germline PGV, were identified by Mutect2 and Varscan2. Locus-specific LOH genomic calls in *TP53* were harmonized with p53 immunohistochemistry (IHC) (below) to determine overall bi-allelic loss status for each LFS-BC and normal breast tissue sample.

### Bioinformatics for Whole Genome Sequencing (WGS)

FASTQ files of 19 LFS-BC WGS were aligned to hg19 using BWA-mem^19^. Sequencing quality control removed three tumors. Somatic copy number variations were called using CNVKit version 0.9.9^22^. A panel of ethnicity and sex matched normal tissues sequenced on the same machine served as a reference set for CNVKit. Using CNVKit copy number segments as input, HRDex was used to determine HRD scores and aneuploidy scores. Copy number burden was calculated as the percentage of bases measured that were in an altered copy number state. Somatic single nucleotide variants were called using Mutect2. A panel of ethnicity and sex matched normals sequenced on the same machine was used to filter out suspected germline variants and technical artifacts. Gnomad was additionally used to filter for common germline variants. TMB was calculated as the number of non-oncogenic SNVs with a depth greater than 20 and an allelic balance greater than 0.05 divided by the number of bases captured and multiplied by 100000. FitMS version 2.3.0^28^ was used to determine the proportion of single nucleotide and copy number variants attributable to known COSMIC mutational signatures.

### Bioinformatics for RNA Sequencing

FASTQ files from 16 LFS-BC, 12 LFS normal breast tissue, 25 non-LFS BC, and 25 non-LFS normal breast tissue were aligned to hg19 using STAR Align version 2.7.8a^29^. Genes were counted using HTSeq-Count version 2.0.3^30^. Differential expression was performed using DESeq2 version 1.38.3^31^. Gene set enrichment analysis (GSEA) was performed using Broad GSEA version 4.2.3^32^. Immune cell type proportions were estimated using CIBERSORTx^33^, MCPCounter^34^ and xCell^35^.

### Analysis of The Cancer Genome Atlas (TCGA) cohorts

To create a TCGA-LFS cohort, carriers of putative germline *TP53* variants were identified from the TCGA pan-cancer analysis^36^. Tumor and matched normal DNA BAM files were downloaded from the Genomic Data Commons (GDC) using a National Center for Biotechnology Information Genotypes and Phenotypes Database (NCBI dbGaP phs000178) approved protocol (#21931) and underwent quality control to ensure the *TP53* variant was found at the expected heterozygous frequency in the normal BAM file and variant classification was performed using the American College of Medical Genetics (ACMG) specific guidelines^37^.

Analysis of TCGA nonLFS-BC whole exome sequencing (WES) was as previously described^17^; briefly, all tumor/normal BAM files were downloaded from GDC and underwent the same variant calling and classification pipeline as above; samples with cancer risk gene PGVs (**Table S4**) were excluded. For the TCGA nonLFS-BC RNAseq analysis, FASTQ files were downloaded from GDC and underwent the same bioinformatics analysis as above. Clinical data was downloaded from cbioportal^38^. TCGA nonLFS-BC were categorized as ER+Her2-, HER2+ and TNBC based on TCGA clinical data and sub-grouped by presence or absence of *TP53* acquired somatic mutation (**Table S5**). The TCGA nonLFS-BC cohort consisted of ER+Her2-(n=263 wild type *TP53* (TP53WT), n=82 mutant *TP53* (TP53Mut)), HER2+ (n=68 TP53WT, n=56 TP53Mut) and TNBC (n=12 TP53WT, n=78 TP53Mut), and n=113 normal breast tissue samples for RNAseq.

### p53 and immune marker immunohistochemistry

For single marker immunohistochemistry (IHC), five-micron sections of formalin-fixed paraffin-embedded tissue were stained using the following anti-human antibodies: DO-7 (p53), C8 (CD8), L26 (CD20), KP1 (CD68) (Dako, Carpinteria, CA); LN10 (CD3) (Leica Microsystems, San Francisco, CA); RM (CD4) (Biocare Medical, Pacheco, CA), 206D (FoxP3) (Biolegend Antibodies, San Diego, CA). Antibody dilutions were as follows: DO-7 (1:60), LN10 (1:1), RM (1:1), C8/144B (1:40), L26 (1:1), KP1 (1:1), 206D (1:100). Staining was performed on a Leica Bond-IIITM instrument using the Bond Polymer Refine Detection DS9800 System (Leica Microsystems, San Francisco, CA. Heat-induced epitope retrieval was done for 20 minutes with ER1 solution (Leica Microsystems, San Francisco, CA). All experiments were done at room temperature. Slides were washed three times between each step with bond wash buffer or water. To quantify the single stain IHC, we used QuPath v0.4.3^39^, on high-resolution IHC brightfield images. We selected the regions of interest within the tissue (normal versus tumor) based on the pathologist marked H&E slide, QuPath algoirthms to identify single cells and quantification of staining intensity and calculation of percent positive cells within each area of tissue.

### Multiplex immunofluorescence (mIF)

Slides for mIF were cut at 5 microns from LFS-BC and non-LFS-BC samples. Each tissue slide was curated by a pathologist to determine if the slide contained tumor tissue. We conducted two independent mIF assays on tumor slides: TIL (Tumor Infiltrating Lymphocytes) and ACT (Activation). The mIF panels were previously standardized for FFPE samples and performed as previously reported^40–42^. TIL assay queried for CD8+, CD4+, CD20+, Cytokeratin (CK), and DAPI. Cancer-cells were determined by CK and DAPI. The ACT assay queried for Granzyme B (GrzmB), Ki67+, CD3+, CK, and DAPI. Granzyme B positivity indicates cytolytic activity and active T-cell mediated killing, Ki67+ is a marker for cell proliferation, and CD3+ tests for the presence of T-cells. A total of 57 samples were tested with the TIL and ACT assays (comprising 7 spontaneous ER+/PR+, 16 spontaneous TNBC, and 33 LFS-BC).

The mIF protocol has been described previously^40–43^. Briefly, FFPE BC sections were submitted to antigen retrieval using EDTA buffer pH=8.0 and incubated with primary/secondary isotype specific antibodies against the targets of interest (e.g., CD3+,CD8+,CD4+ etc.). Secondary antibodies were then labeled using specific fluorophores and signal was amplified using dextran polymers and/or tyramide-based conjugates. The stained sections were acquired using a multispectral microscope station including several 20X magnification fields of view (FOVs) (median 78 FOVs, mean 83 +/-53). We partitioned the FOVs into tumor area, stroma, and surrounding normal tissue. All examined slides were first vetted to include at least 5% area including tumor tissue (median 74 tumor fields of view, mean 81+/-54 tumor fields of view). The spatially resolved signal quantitation for the mIF markers was conducted as described^40,43^ using the AQUA® platform based on marker co-localization; and the InForm® software based on single-cell segmentation and phenotyping. As each tissue section had variable tumor content and to assess the possible impact of spatial immune-cell heterogeneity, we then examined the top 10% FOVs for each slide, which represented the highest marker level or immune-cell density within that slide. Each FOV was quantitatively described by marker signal level in marker-defined compartment (AU), spatial information (X and Y coordinates) and marker-positive cell density (per mm^2^).

### Statistical analysis

Genomics statistical analyses were completed in R. Comparison of continuous variables were completed using a two-tailed student’s t test. Comparison of alteration rates was done using Fisher’s exact or Chi squred tests. For the mIF analyses, statistical comparisons were conducted with the Mann-Whitney U-tests. Linear and rank correlations were conducted with the Pearson and Spearman correlations. Spatial heterogeneity was computed utilizing the Rao entropy^43^.

## Acknowledgements

Research support for this study was generously provided by The National Institutes of Health (K08CA215312, KNM); Li Fraumeni Syndrome Association (AJL, KNM); Burroughs Wellcome Foundation (1017184, KNM); Basser Center for BRCA at the University of Pennsylvania (MT, RH, KNM) NIH/NCI Cancer Center Support Grant P30 CA008748 (NIH/NCI Cancer Center Support Grant P30 CA008748 and an ASPIRE award from the Mark Foundation BG, DH); ASPIRE award from the Mark Foundation (BG,DH).

