## Supplementary Figures for "Distinct genomic and immunologic tumor evolution in germline *TP53-*driven breast cancers"

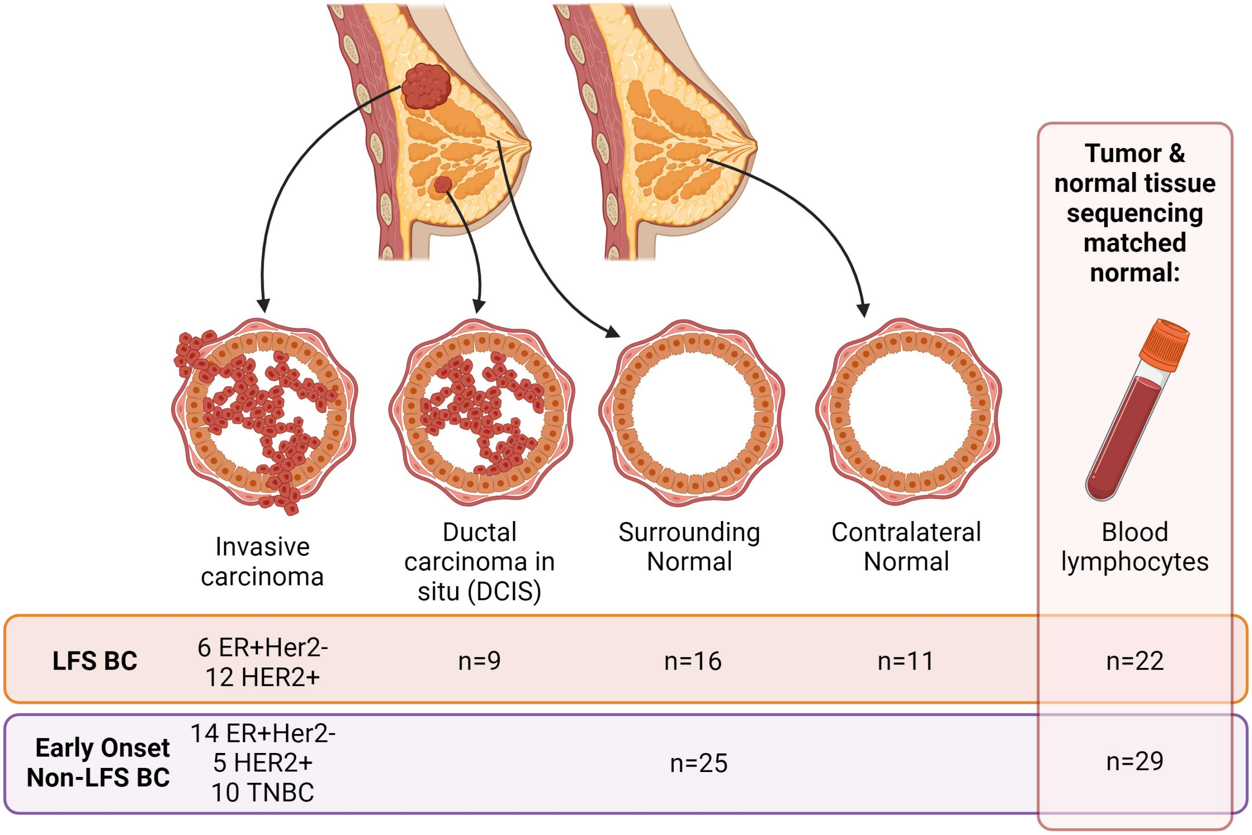


**Figure S1: Tissues analyzed in this study from Penn Medicine.** Breast tumors including both invasive ductal carcinoma (IDC) and ductal carcinoma in situ (DCIS), surrounding normal breast tissue, contralateral normal breast tissue, and blood specimens were obtained with informed consent via a Penn/CHOP IRB approved protocol (LFS/TP53 Biobank) for patients with Li Fraumeni Syndrome (LFS) or the Penn Medicine Biobank (PMBB) for early-onset non-LFS patients. These tissues were subjected to sequencing, transcriptomic and immunohistochemistry analyses. The LFS-BC cohort contained six ER+Her2- IDC, nine HER2+ IDC, nine DCIS, 16 surrounding normal, 11 contralateral normal tissue samples from 21 patients; blood specimens were available from all 22 patients. The early onset non-LFS BC cohort contained 14 ER+Her2- BC, five HER2+ BC and 10 TNBC, and 25 surrounding normal tissue samples from 29 patients with matched normal blood samples. BC, breast cancer; ER: Estrogen Receptor; HER2: receptor tyrosine-protein kinase erbB-2; LFS, Li Fraumeni Syndrome; TNBC: triple negative breast cancer.

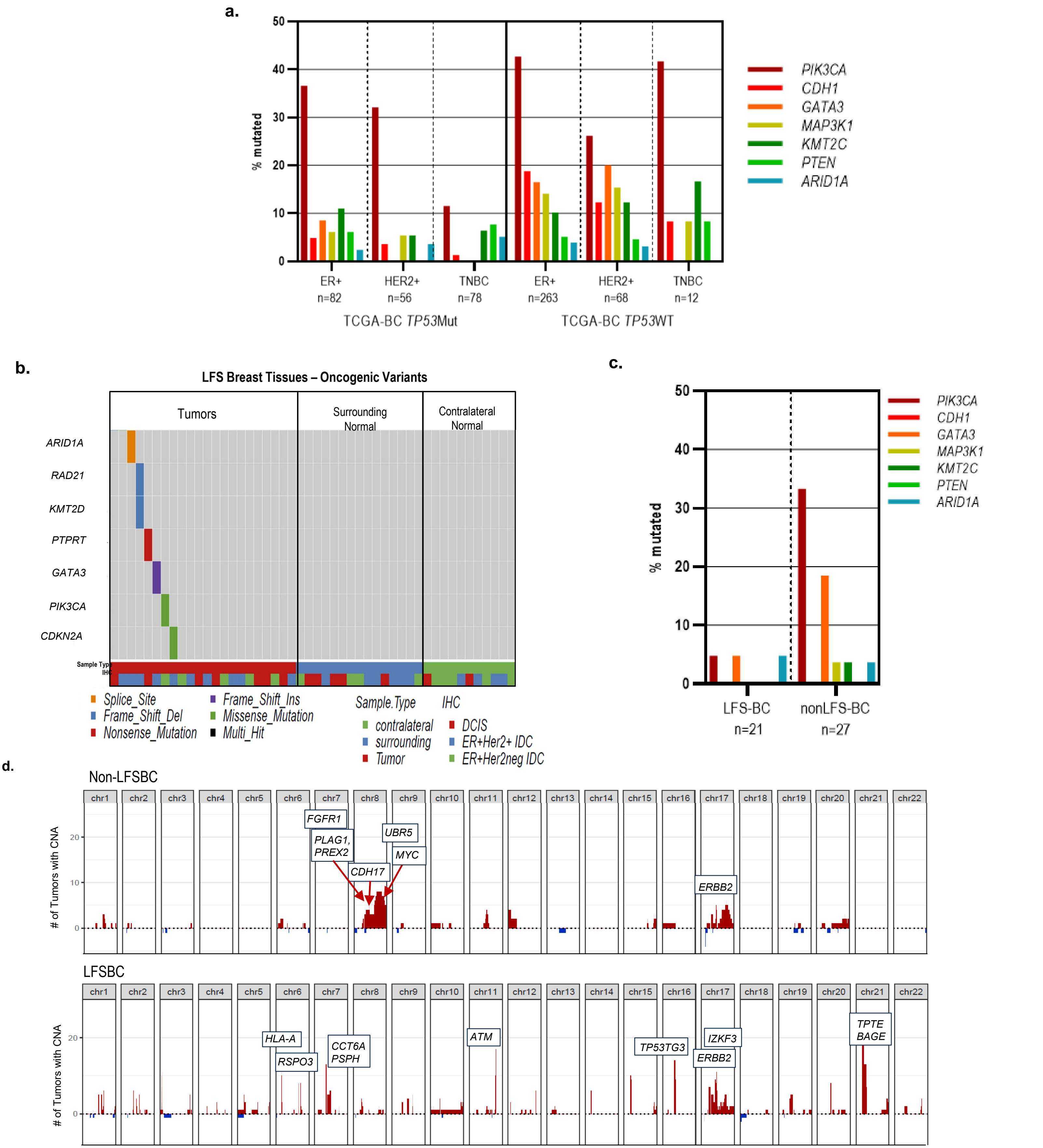


**Figure S2: Rates of common breast cancer oncogenic alterations stratified by p53 status. a.** Fraction of tumors from TCGA-BC cohort with oncogenic mutations in the indicated genes. Tumors are stratified by somatic *TP53* mutation status. **B.** Common breast cancer oncogenic alterations identified by Mutect in LFS-BC and normal tissues. **c.** Fraction of LFS-BC and early onset nonLFS-BC with oncogenic mutations in the indicated genes. **d.** Chromosomal plot of amplifications in nonLFS-BC and LFS-BC. Callouts represent oncogenes from Cancer Gene Census. Del, deletion; DCIS, ductal carcinoma in situ; ER, estrogen receptor; HER2, erbb2 receptor; IDC, invasive ductal carcinoma; LFS-BC, Li Fraumeni Syndrome breast cancer; Ins, insertion; TNBC, triple negative breast cancer; *TP53*Mut, positive for somatic *TP53* mutation; *TP53*WT, negative for somatic *TP53* mutation.

**a.**

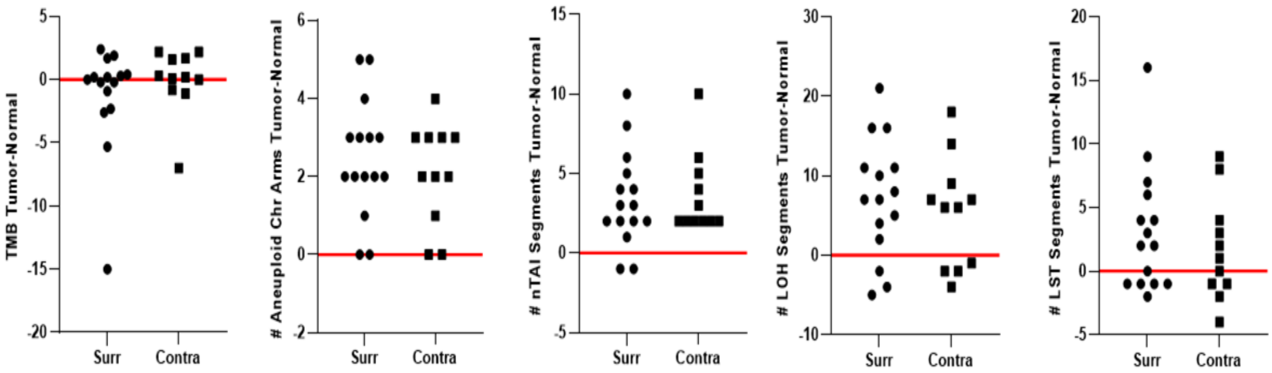


**b.**

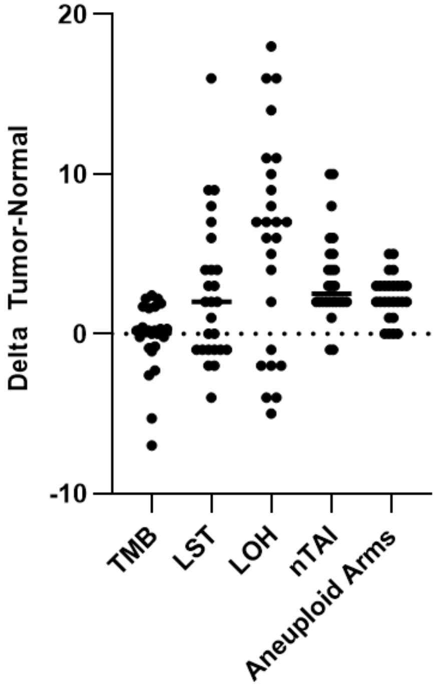


| Tumor - Normal Delta (26 comparisons) | # tumor enriched (delta>1) | frac tumor enriched |
|---|---|---|
| Mutations | 7 | 27% |
| LOH segments | 19 | 73% |
| LST segments | 17 | 65% |
| nTAI segments | 24 | 92% |
| Aneuploid chr arms | 26 | 100% |

**Figure S3: Genomic instability in pairs of LFS-BC versus LFS normal breast tissues. a.** Ratio of TMB, number of aneuploid (Chr) arms, number of nTAI segments, number of genomic LOH segments and number of LST segments in individual LFS-BC sample versus its own matched surrounding (Surr, circles) or contralateral (Contra, squares) normal breast tissue sample. Ratios above the red line (0) indicate the genomic instability measure was higher in the tumor than the normal breast tissue. Genomic instability measures were derived from matched analyses of tumor versus the patient's blood data and normal breast tissue data versus the patient's blood data. Each data point represents one tumor-normal pair. **b.** Delta of the TMB, LST, LOH, nTAI, aneuploid arms measures comparing individual LFS-BC sample versus its own normal (26 comparisons) and fraction of tumors with enrichment of the genomic instability measure. Chr, chromosome; LFS-BC, Li Fraumeni Syndrome breast cancer; LOH, loss of heterozygosity; LST, large state transitions; nTAI, non-telomeric allelic imbalance; TMB, tumor mutation burden.

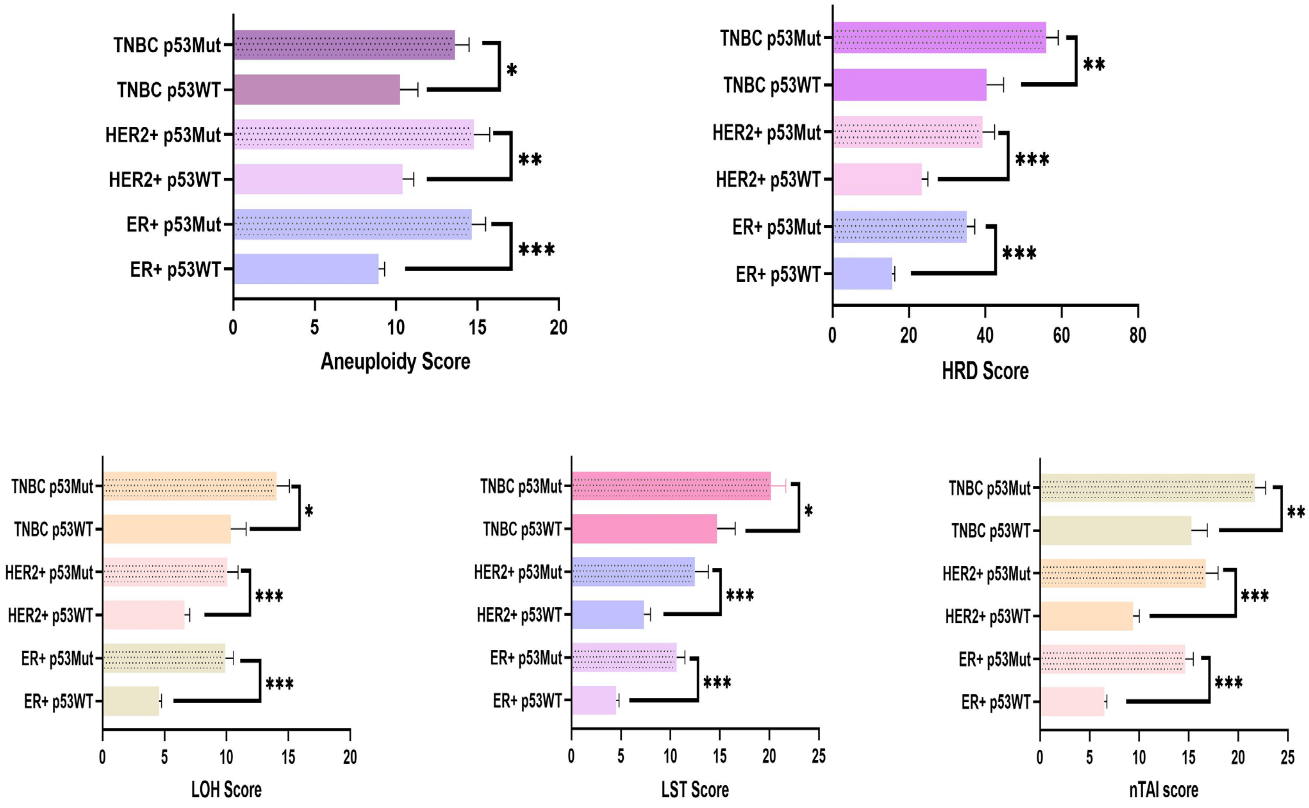


**Figure S4: Chromosomal instability stratified by somatic *TP53* mutation status in TCGA-BC.** Average aneuploidy score, average HRD score, and average number of segments of genomic LOH, LST, and nTAI in ER+Her2-, HER2+ and TNBC from TCGA with or without a somatic *TP53* mutation. ER: Estrogen Receptor; HER2: receptor tyrosine-protein kinase erbB-2; HRD, homologous recombination score; LOH, loss of heterozygosity; LST, large state transitions; nTAI, non-telomeric allelic imbalance; TNBC: triple negative breast cancer.

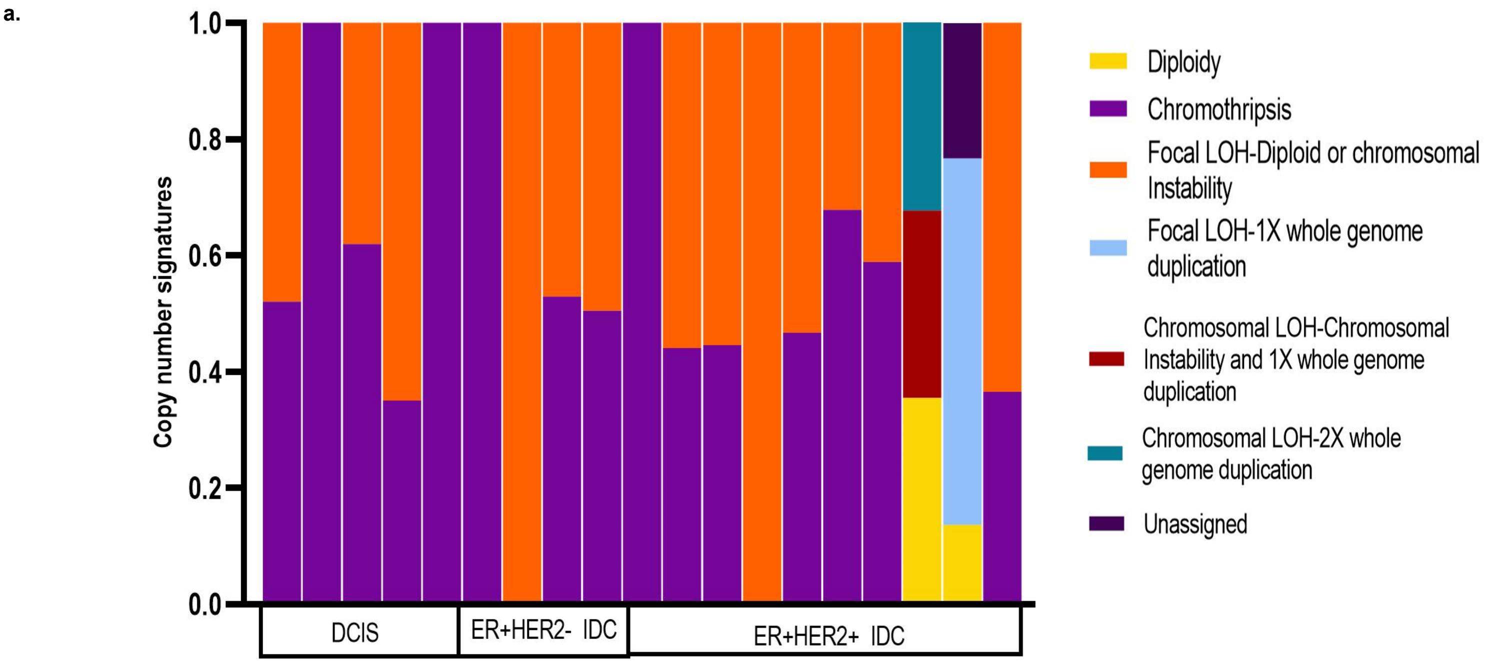


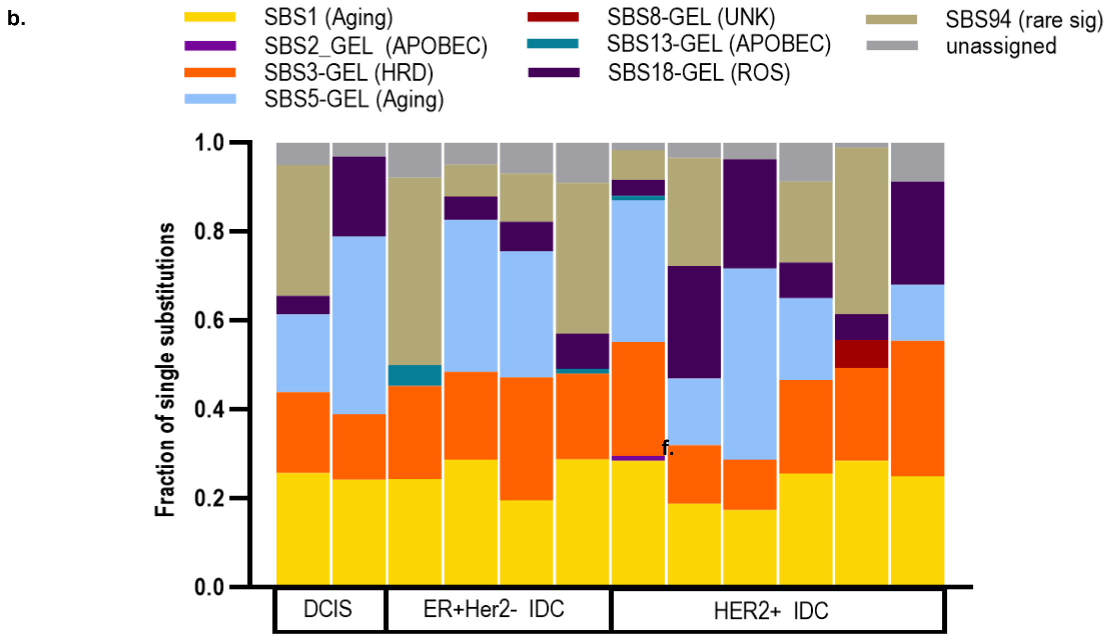


**Figure S5: Mutational processed in LFS-BC.** (a) Copy number signature profile of LFS-BC from whole genome sequencing. (b) Single base pair mutational signature profiles derived from FitMS of LFS-BC by whole genome sequencing. LOH, loss of heterozygosity; HRD, homologous recombination deficiency; ROS, reactive oxygen species/ SBS, single base substitution; UNK, unknown

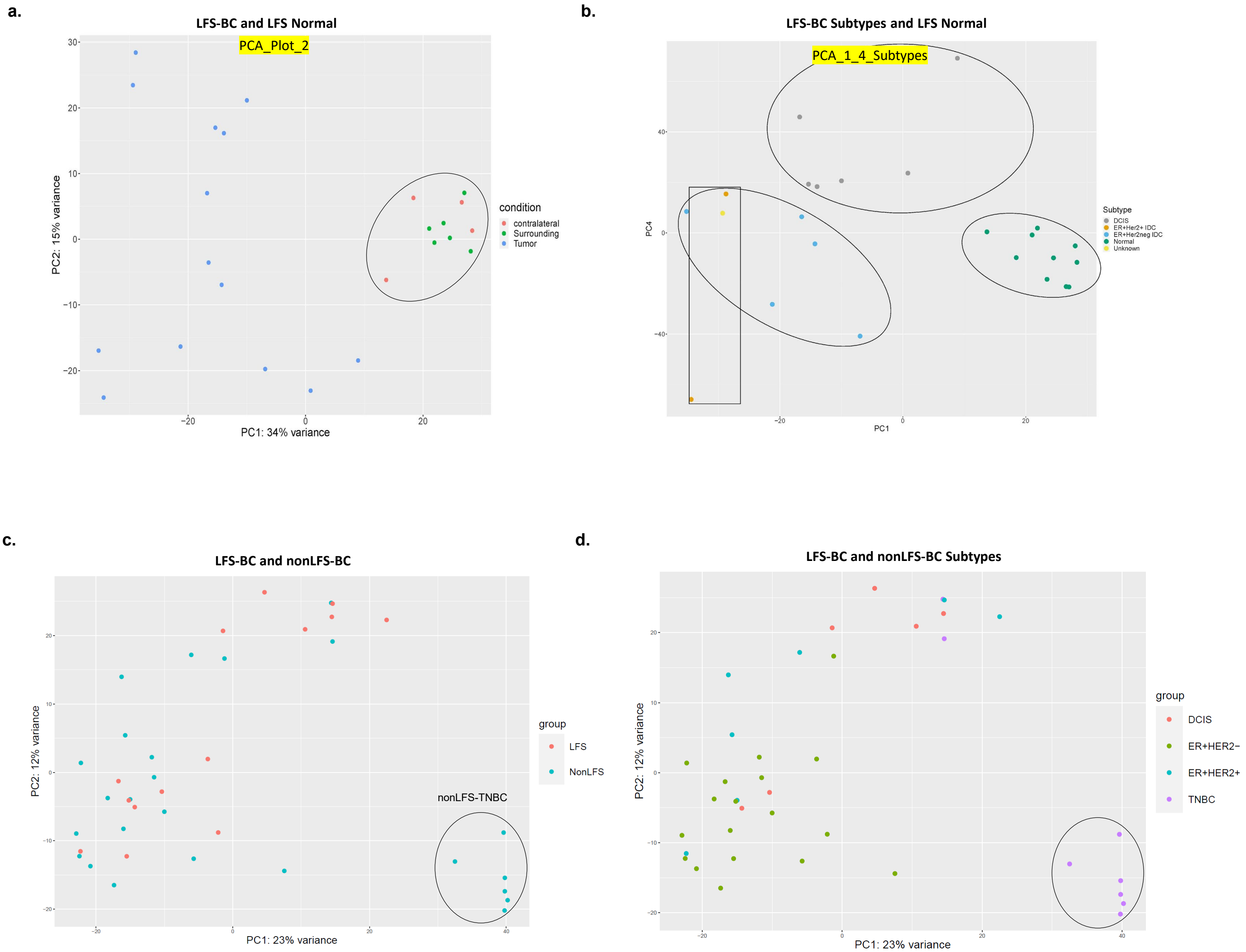


**Figure S6: PC plots showing expression data subgroupings**. **a.** PC plot of RNAseq data from LFS-BC and LFS normal breast tissue showing clustering of tumors separate from both contralateral and surrounding normal breast tissue. **b.** PC plot of RNAseq data from LFS-BC stratified by subtype (DCIS, ER+Her2- and HER2+ IDC) and LFS normal breast tissue showing clustering of DCIS separate from invasive tumors and separate from normal breast tissue. **c.** PC plot of RNAseq data from LFS-BC and non-LFSBC showing intermixing of most tumors except one grouping (circle). **d.** PC plot of RNAseq data from LFS-BC and nonLFS-BC stratified by subtype (DCIS, ER+Her2-, HER2+ IDC and TNBC) showing clustering of TNBC tumors separate from invasive tumors and separate from normal breast tissue. BC, breast cancer; DCIS, ductal carcinoma *in situ*; ER: Estrogen Receptor; HER2: receptor tyrosine-protein kinase erbB-2; IDC, invasive ductal carcinoma; LFS, Li Fraumeni Syndrome; TNBC: triple negative breast cancer.

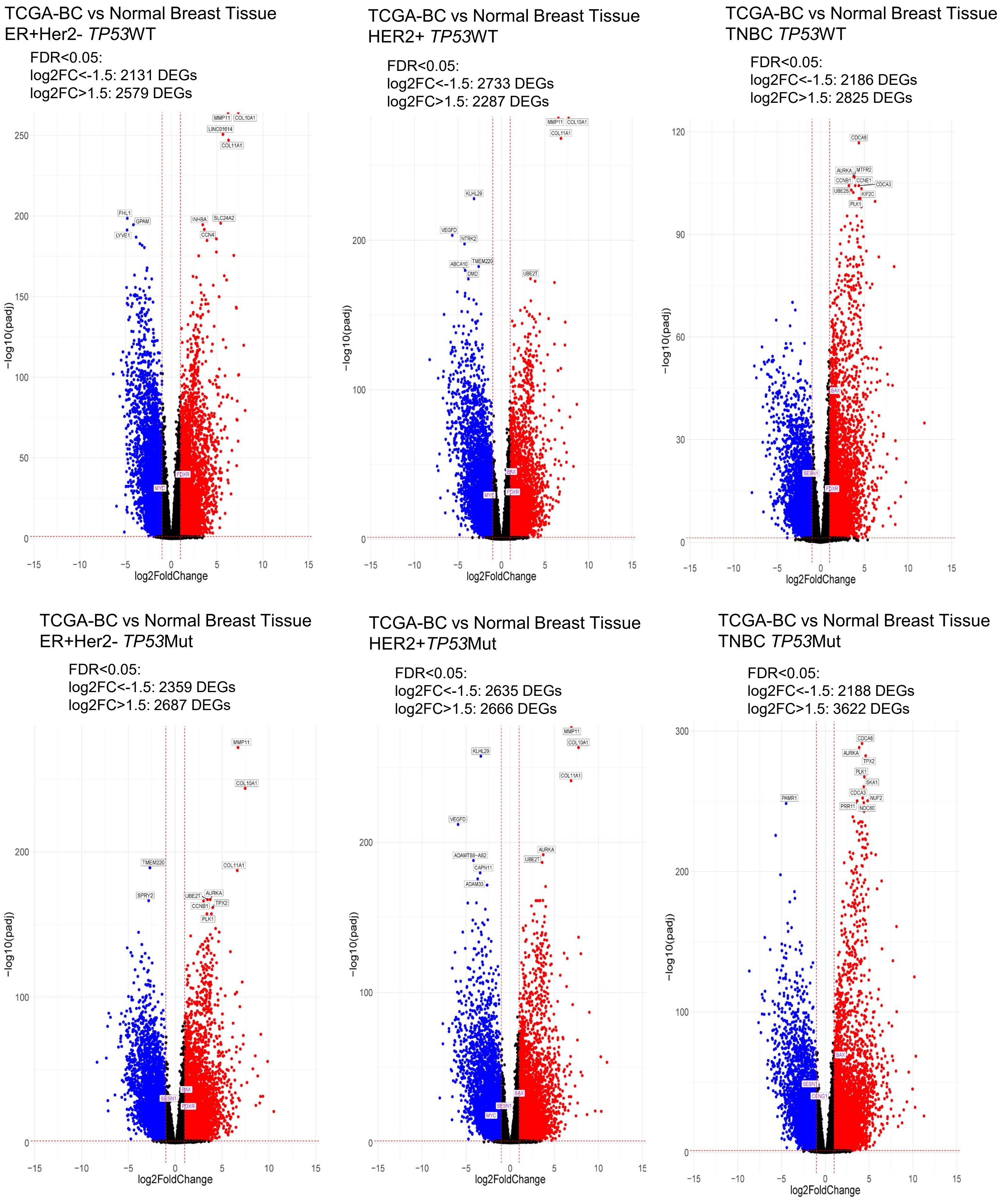


**Figure S7: Differential gene expression analysis in TCGA-BC cohort stratified by somatic *TP53* mutation status and hormone receptor subtype.** ER: Estrogen Receptor; HER2: receptor tyrosine-protein kinase erbB-2; TNBC: triple negative breast cancer (Blue: Down, Red: Up and Black: No change)

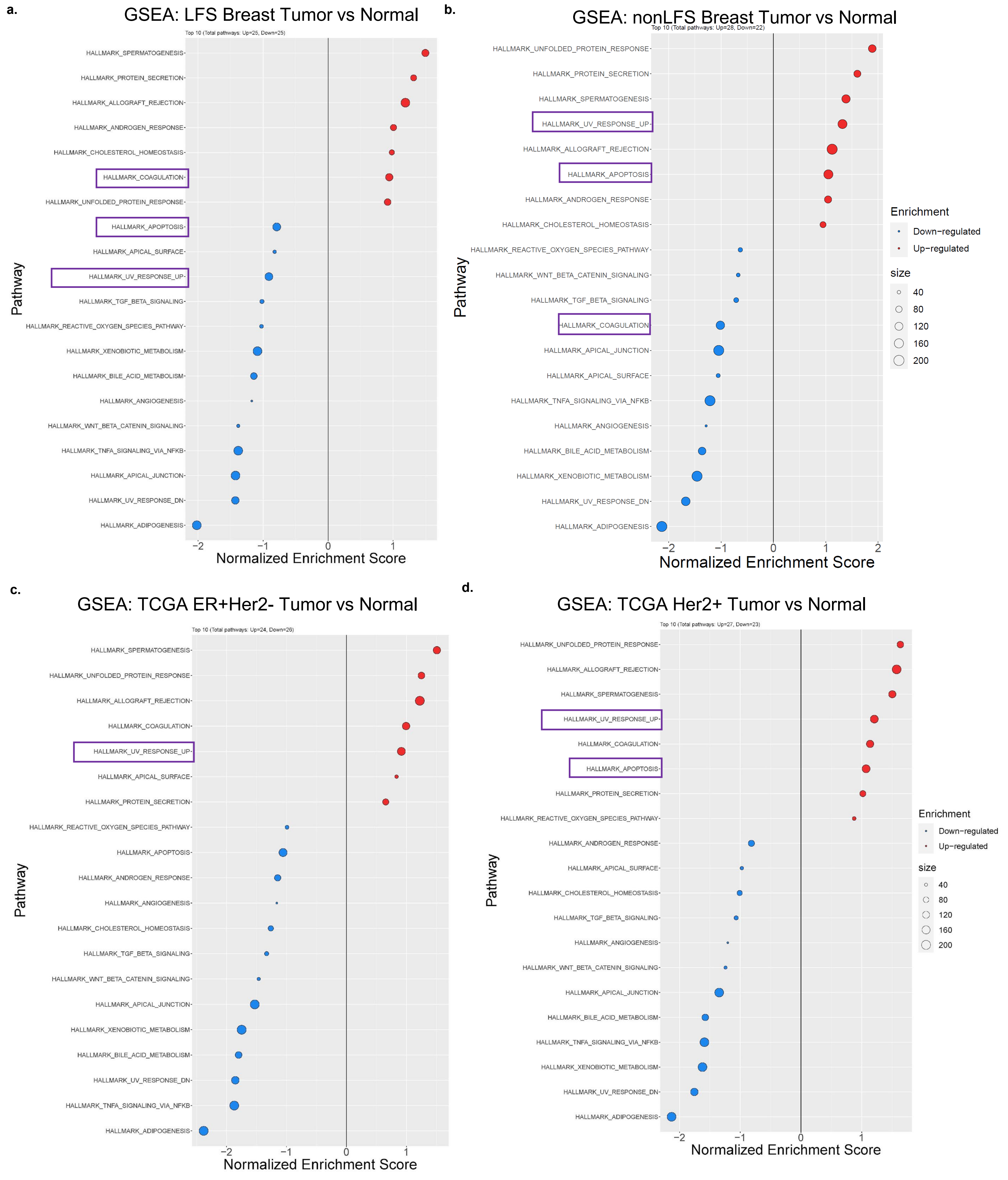


**Figure S8: Gene set enrichment analysis (GSEA) of LFS-BC and non-LFS BC**. **a.** GSEA of LFS-BC compared to LFS normal breast tissue. **b.** GSEA of nonLFS-BC (ER+) compared to nonLFS normal breast tissue. **c.** GSEA ER+Her2- nonLFS-BC compared to normal breast tissue from TCGA. **d.** GSEA Her2+ nonLFS-BC compared to normal breast tissue from TCGA. BC: Breast cancer; ER: Estrogen Receptor; HER2: receptor tyrosine-protein kinase erbB-2; LFS: Li Fraumeni Syndrome.

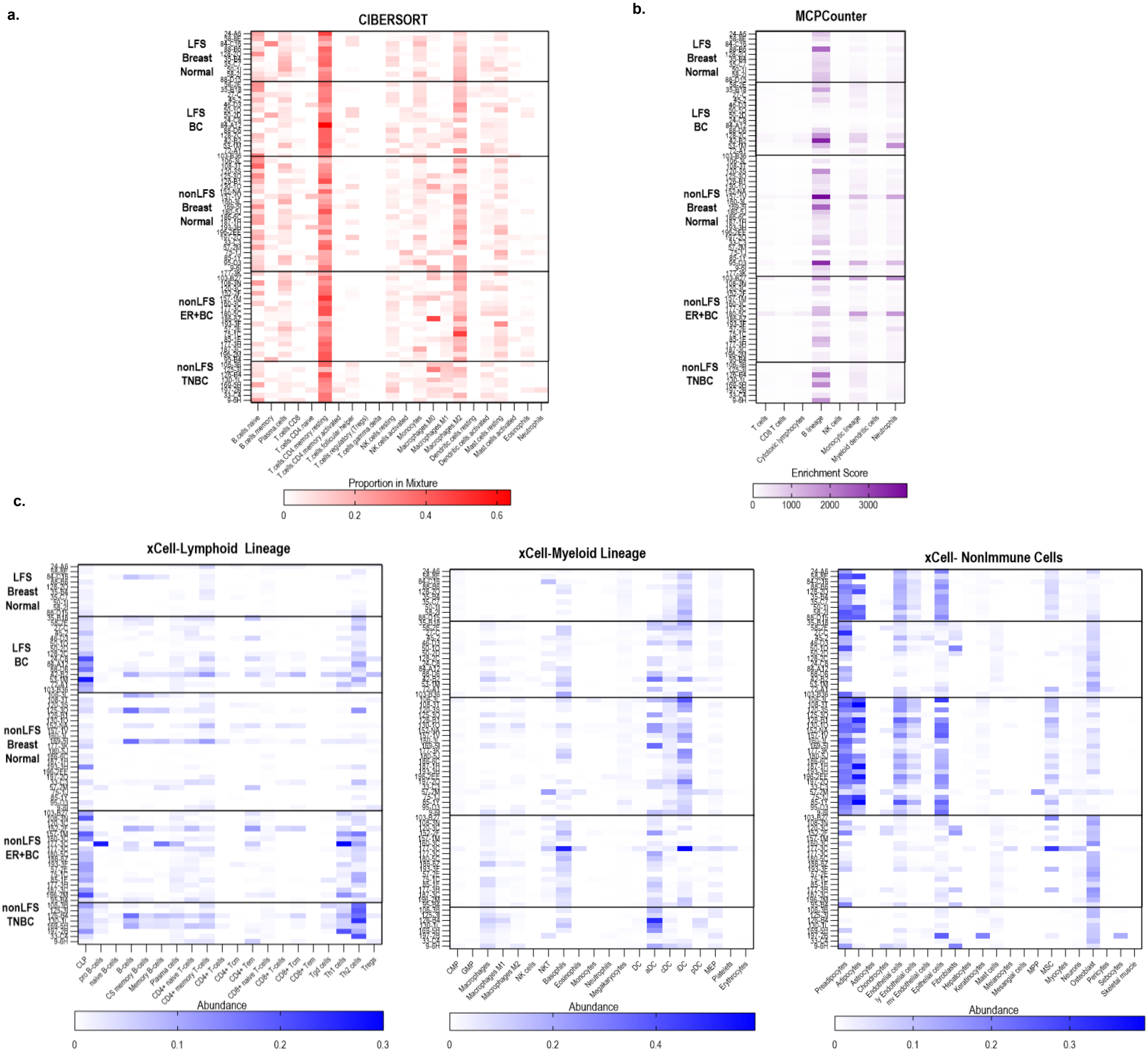


**Figure S9: Analysis of tumor microenvironment . a.** CIBERSORT analysis of RNAseq data estimating immune cell proportions in LFS and nonLFS-BC and normal breast tissues. **b.** MCPCounter analysis of RNASeq data estimating enrichments scores of immune cell proportions in LFS and nonLFS-BC and normal breast tissues. **c.** xCell analysis of RNASeq data estimating abundance of lymphoid lineage immune cells, myeloid lineage immune cells and non-immune cells in LFS and nonLFS-BC and normal breast tissues. Breast cancer; ER: Estrogen Receptor; HER2: receptor tyrosine-protein kinase erbB-2; LFS: Li Fraumeni Syndrome; TNBC: Triple negative breast cancer.

a.

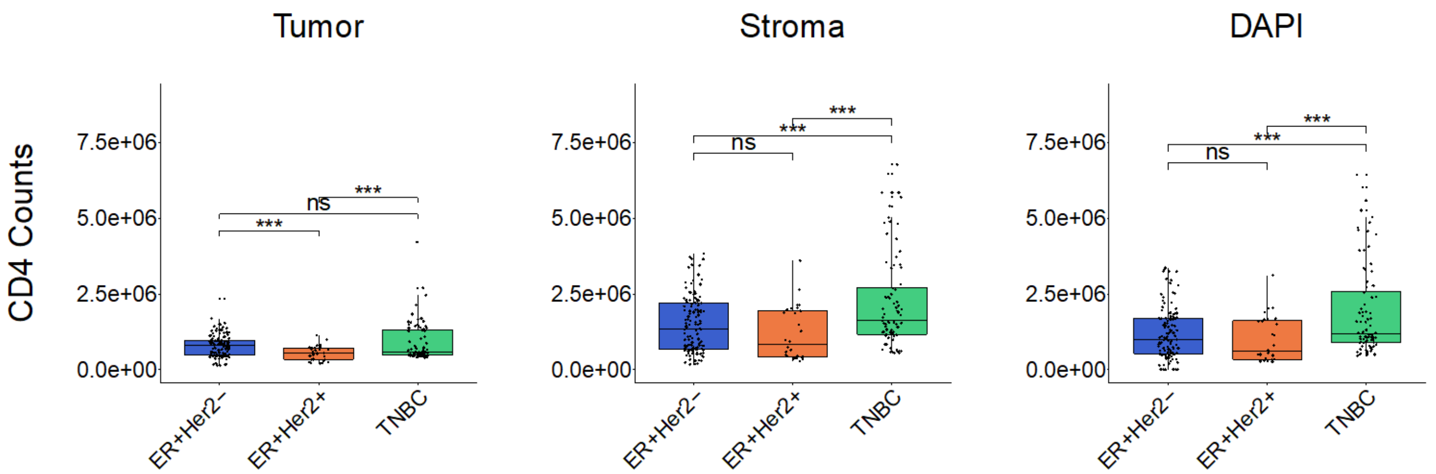


b.

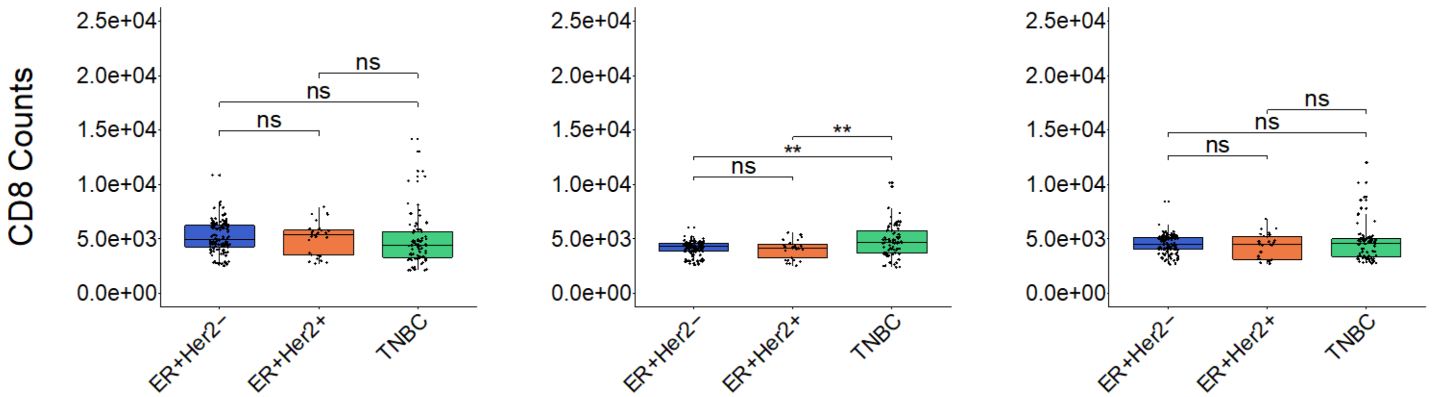


c.

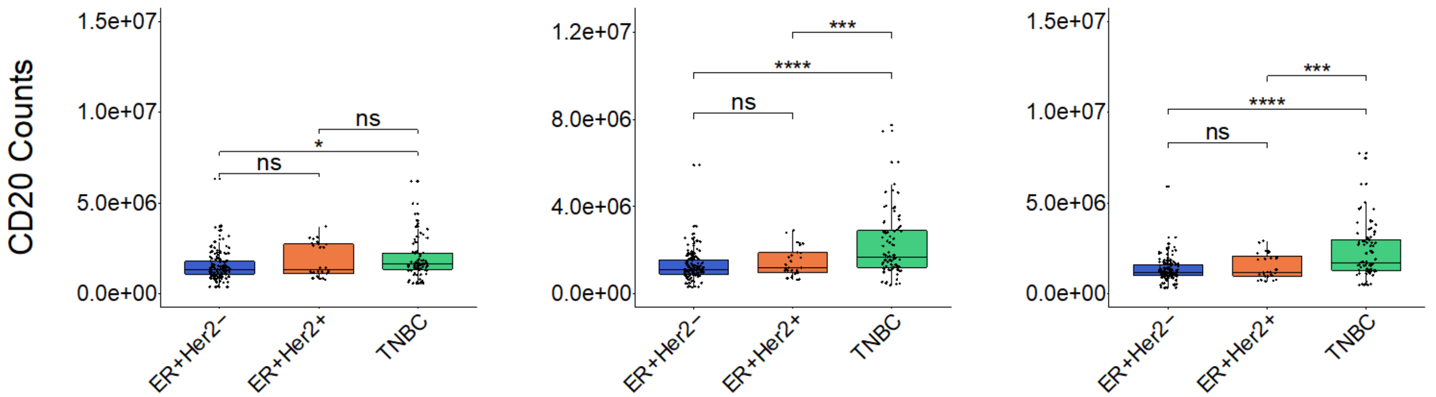


**Figure S10: Immune infiltration in ER+Her2-BC, HER2+-BC and TNBC in premenopausal nonLFS-BC**. **a.** CD4+ cells in tumor, stromal and DAPI channel using multiplex immunohistochemistry. **b.** CD8+ cells in tumor, stromal and DAPI channel using multiplex immunohistochemistry. **c.** CD20+ cells in tumor, stromal and DAPI channel using multiplex immunohistochemistry. ER: Estrogen Receptor; HER2: receptor tyrosine-protein kinase erbB-2; LFS: Li Fraumeni Syndrome; TNBC: Triple negative breast cancer.

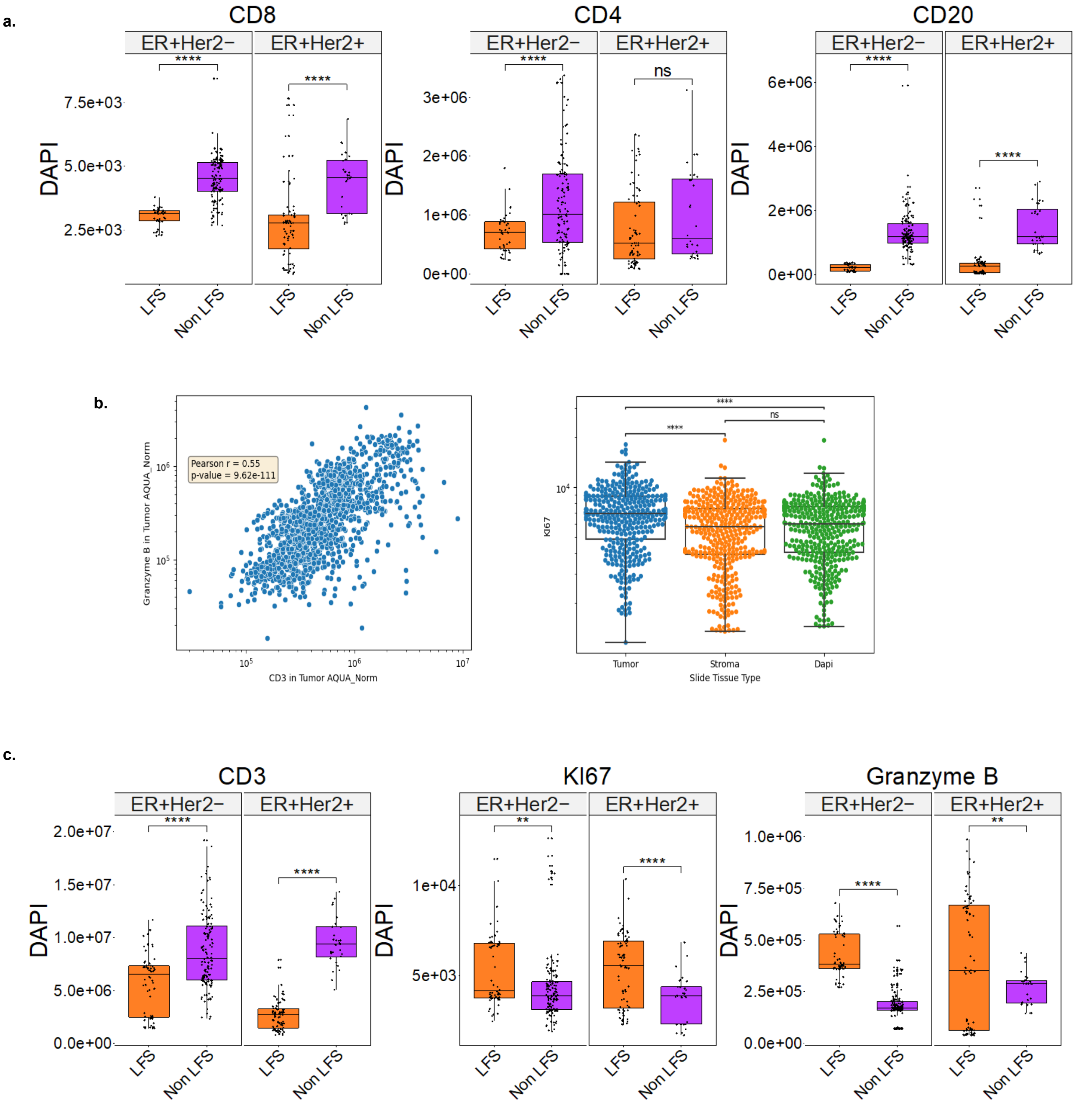


**Figure S11: Multiplex Immunohistochemistry to study the tumor immune microenvironment in LFS-BC versus nonLFS-BC. a.** Levels of CD8+, CD4+ and CD20+ cells in the DAPI compartment in LFS-BC versus nonLFS-BC. **b.** Correlation of Granyzme B and CD3 staining in tumor and demonstration of higher Ki67 staining in tumor versus stroma and DAPI. **c.** T-cell activation panel in LFS-DCIS compared to LFS-IDC. ACT, T-cell activation panel; BC, breast cancer; CK, cytokeratin; ER: Estrogen Receptor; GrzmB, granzyme B; LFS: Li Fraumeni Syndrome; TNBC: Triple negative breast cancer.

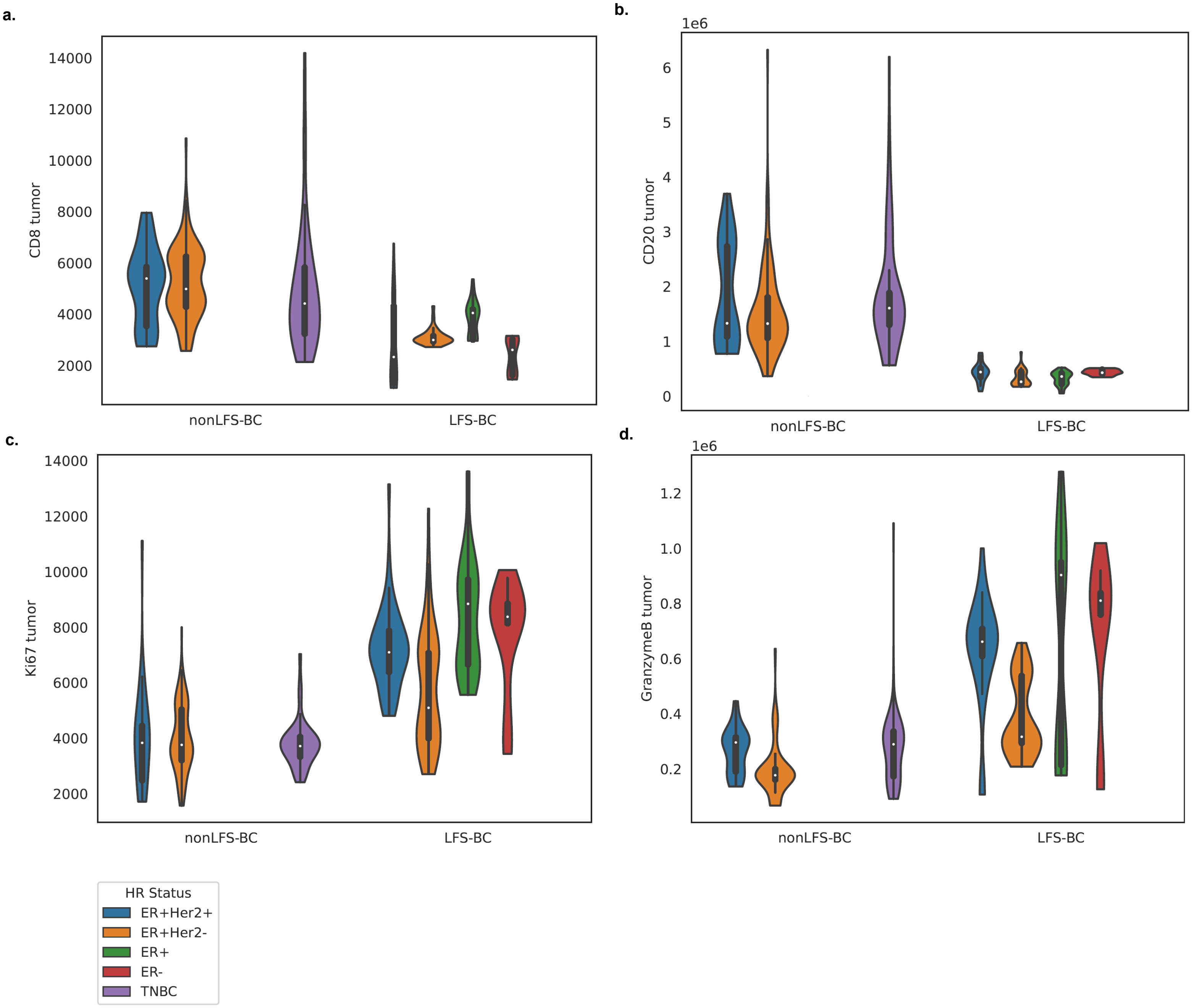


**Figure S12: Multiplex immunohistochemistry analysis in expanded cohort of LFS-BC versus nonLFS-BC. a.** Tumor infiltration of CD8+ cells in HER2+ BC, ER+Her2- BC, ER+ DCIS, ER- DCIS, and TNBC in nonLFS-BC and LFS-BC from both pre-menopausal and post-menopausal patients. **b.** Tumor infiltration of CD20+ cells in HER2+ BC, ER+Her2- BC, ER+ DCIS, ER- DCIS, and TNBC in nonLFS-BC and LFS-BC from both pre-menopausal and post-menopausal patients. **c.** Tumor infiltration of Ki67+ cells in HER2+ BC, ER+Her2- BC, ER+ DCIS, ER- DCIS, and TNBC in nonLFS-BC and LFS-BC from both pre-menopausal and post-menopausal patients. **d.** Tumor infiltration of GranzymeB+ cells in HER2+ BC, ER+Her2- BC, ER+ DCIS, ER- DCIS, and TNBC in nonLFS-BC and LFS-BC from both pre-menopausal and post-menopausal patients. DCIS, ductal carcinoma in situ; ER: Estrogen Receptor; HER2: receptor tyrosine-protein kinase erbB-2; LFS: Li Fraumeni Syndrome; TNBC: Triple negative breast cancer.

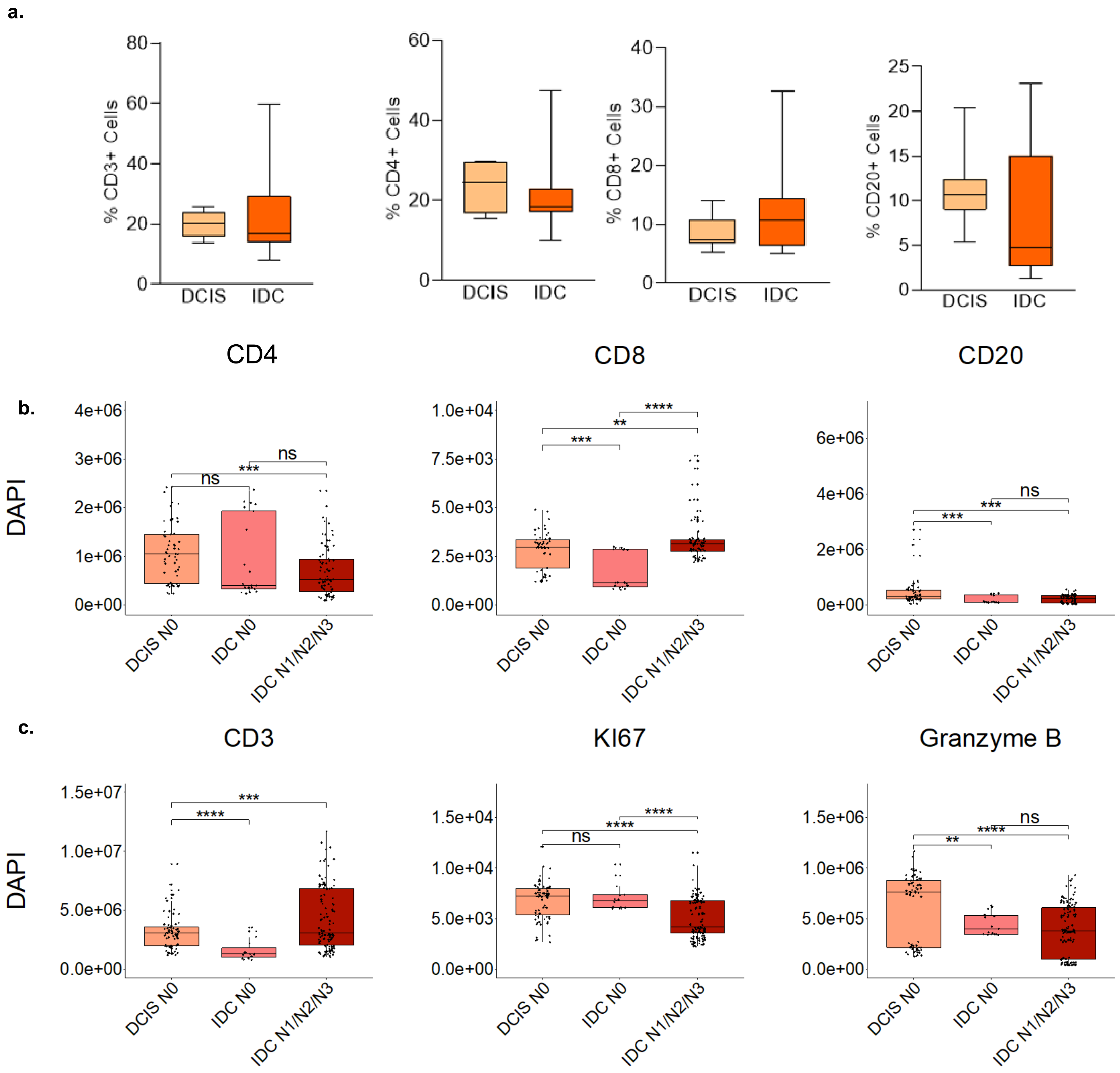


**Figure S13: Immune staining in LFS-BC stratified by DCIS and IDC.** **a.** Single immune staining in LFS DCIS versus LFS IDC for CD3+ T cells, CD4+ T cell, CD8+ T cells and CD20+ T cells. **b.** Levels of CD8+, CD4+ and CD20+ cells in the DAPI compartment in LFS-DCIS vs LFS-IDC. **c.** Levels of CD3+, granzymeB and Ki67+ cells in the DAPI compartment in LFS-DCIS vs LFS-IDC. DCIS, ductal carcinoma in situ; IDC, invasive ductal carcinoma; LFS, Li Fraumeni Syndrome.
